# Visual object topographic motifs emerge from self-organization of a unified representational space

**DOI:** 10.1101/2022.09.06.506403

**Authors:** Fenil R. Doshi, Talia Konkle

## Abstract

The object-responsive cortex of the visual system has a highly systematic topography, with a macro-scale organization related to animacy and the real-world size of objects, and embedded meso-scale regions with strong selectivity for a handful of object categories. Here, we use self-organizing principles to learn a topographic representation of the data manifold of a deep neural network representational space. We find that a smooth mapping of this representational space showed many brain-like motifs, with (i) large-scale organization of animate vs. inanimate and big vs. small response preferences, supported by (ii) feature tuning related to textural and coarse form information, with (iii) naturally emerging face- and scene-selective regions embedded in this larger-scale organization. While some theories of the object-selective cortex posit that these differently tuned regions of the brain reflect a collection of distinctly specified functional modules, the present work provides computational support for an alternate hypothesis that the tuning and topography of the object-selective cortex reflects a smooth mapping of a unified representational space.

---

Extensive empirical research has charted the spatial layout of tuning preferences along the ventral visual stream (occipitotemporal cortex “OTC” in humans and inferior temporal “IT” cortex in monkeys; for review see: Grill-Spector & Weiner, 2014; Tsao et al., 2006; Freiwald & Tsao, 2010; Ungerleider, 1982; Kanwisher, 2010; Ungerleider and Bell, 2011). At a macro-scale, there are two major object dimensions which have been shown to elicit systematic large-scale response topographies, related to the distinction between animate and inanimate objects (Grill-Spector & Weiner, 2014; Haxby et al., 2011; Chao et al., 1999; Martin et al., 1996; Mahon & Caramazza, 2011; Naselaris et al. 2012, Sha et al. 2015; Martin, 2007) and the distinction between objects of different real-world sizes (Konkle & Oliva, 2012; Konkle & Caramazza, 2013; Julian et al., 2017). Further research has shown that these seemingly high-level animacy and object size distinctions are in fact primarily accounted for by differences in tuning along more primitive visuo-statistical features that meaningfully co-vary with these high-level properties (e.g. at the level of localized texture and coarse form information; Long et al., 2018, Jagadeesh & Gardner, 2022, Coggan et al., 2022).

At a meso-scale, there is a hallmark mosaic of category-selective regions scattered across this cortex, defined by their spatially clustered and highly selective responses to a particular category--e.g., faces, bodies, letters, and scenes (Kanwisher et al., 1997; McCarthy et al., 1997; Tsao et al., 2006; Downing et al., 2001; Peelen & Downing, 2005; Epstein & Kanwisher, 1998; Aguirre et al., 1998; Nasr et al., 2011; Kanwisher, 2010; Puce et al., 1996; Polk et al., 2002)--with no such highly selective regions for other categories like cars and shoes (Downing et al., 2006). Initially, it was unclear whether these regions should be considered “stand-alone modules” which are unrelated to the object tuning preferences of the surrounding regions (Op de Beeck et al., 2008). However, it is increasingly clear that there is a systematic encompassing structure in the cortical organization, where the face, body, and scene-selective regions fall systematically and meaningfully within this larger-scale animacy and object size organization (Konkle & Caramazza, 2013; Grill-Spector & Weiner, 2014; Bao et al., 2020). This systematic map of object tuning, at both macro- and meso-scales, has led to an extensive debate and discussion--why are these macro and meso-scale object distinctions evident and not others, and why are they spatially organized this way (e.g. Malach et al. 2002; Kanwisher, 2010; Mahon & Caramazza, 2011; Konkle & Oliva, 2012; Grill-Spector & Weiner, 2014; Op de Beeck et al., 2019; Conway, 2018)?

On one theoretical account, the tuning and topography of neurons in the object-selective cortex could be conceived of as jointly capturing a unified representational space, which is smoothly mapped along the cortical surface (Bao et al., 2020; Konkle & Caramazza, 2013). That is, the *tuning* of this entire population of neurons is best understood together, as part of an integrated, large-scale population code, with features designed to discriminate *all* kinds of visual input, including faces (Ishai et al., 1999; Haxby et al., 2001; Haxby et al., 2011; Chao et al., 1999; Dicarlo & Cox, 2007). Further, this account continues that this multi-dimensional representational space is mapped along the 2-dimensional cortex such that similar tuning is nearby, and more distinct tuning is farther apart (Behrmann & Plaut, 2013; Plaut & Behrman, 2011; Grill-Spector & Weiner, 2014; Cowell and Cotrell, 2013). On this account, animacy and object size distinctions have a large-scale organization because they are related to the major dimensions of this unified visual feature space. At the same time, meso-scale regions for faces, bodies, and scenes emerge due to their related visuo-statistical characteristics with other object categories, without requiring other specialized mechanisms.

This theoretical account of the tuning and topography of the object-selective cortex has been challenging to test, as there were no image-computable feature spaces rich enough to categorize many kinds of objects (Kourtzi & Conner, 2011). However, deep neural networks trained to do many-way object categorization, without any special feature branches set aside for some categories, provide precisely this kind of a unified representational space (Prince & Konkle, 2020; Khosla & Wehbe, 2022). Indeed, recently, Bao et al., (2020) used a late layer of a deep neural network (AlexNet) to operationalize such a unified representational space, proposing that the monkey IT organization can be thought of as a coarse map of this space. In so doing, they could predict the tuning of previously uncharted regions of the primate visual cortex based on the major dimensions of the deep neural network feature space, and they linked animacy and object protrusion distinctions to the major principal components of this DNN space. Relatedly, Huang et al., (2022) have found that information about the real-world size of objects is encoded along the second principal component of the late stages of deep neural networks. Further, Vinken et al., 2022 recently demonstrated that face-selective neurons in IT could be accounted for by the feature tuning learned in these same object-trained deep neural networks (also see Prince & Konkle, 2020; Murty et al., 2021; Khosla & Wehbe, 2022). Thus, deep neural networks clearly operationalize a multi-dimensional representational encoding space that has information about these well-studied object distinctions.

One critical missing component of this theoretical account, though, is how to bridge from the multi-dimensional representational spaces of deep neural networks to the spatialized tuning of the cortical sheet—that is, to have a computational account of not only what the tuning is, but also where it is located on a two-dimensional surface. Concurrently, a variety of approaches are emerging to bring spatial organization in deep neural networks, all of which operate at different levels of abstraction regarding the underlying mechanisms (Lee et al., 2020; Blauch et al., 2022; Zhang et al., 2021; Keller et al., 2021). Here, we cast the problem of topography as one of data-manifold mapping, leveraging Kohonen self-organizing maps (Kohonen, 1990). This computational approach aims to reveal the similarity structure of natural images implicit in the deep neural network feature space, by smoothly embedding a two-dimensional sheet into the multi-dimensional feature space to capture this structure. This computational approach has previously been successfully used to account for other representational-topographic signatures found along the cortex, including the large-scale multiple-mirrored map topography of the early visual system areas (Konkle, 2021; see also Durbin & Mitchison 1990; Obermayer et al., 1990), the large-scale body-part and action topography of the somatomotor cortex (Aflano & Graziano, 2006; Graziano & Aflano, 2007; Aflano & Graziano, 2011), and even early explorations of object category topography (Cowell & Cottrell, 2013).

We developed a framework to train a self-organizing map (SOM) over the feature space learned in the late stage of a deep neural network model, and then probed for several key signatures of the ventral stream topography. Doing so revealed several brain-like macro- and meso-scale response topographies, which naturally emerge from a smooth mapping of the DNN feature space, including the formation of localized category-selective regions for faces and scenes. However, not all known topographic signatures of the ventral visual pathway were evident in the modeled topography. Broadly, this work provides computational plausibility for a theoretical account in which the organization of object-selective cortex can be understood as a smooth mapping of a unified representational space along a two-dimensional sheet. Further, under these assumptions, the departures between the object representation in DNNs and the human brain reveal clear modeling directions to drive towards a more brain-like representational system.

## RESULTS

### Self-organizing maps learn the data manifold of a deep neural network feature space

Here, we use a standard pre-trained Alexnet neural network (Krizhevsky et al., 2012), focusing on the representation of natural images in the penultimate layer (relu7) prior to the output layer. This stage reflects the most transformed representational format from the pixel-level representation. Within this layer, the set of natural images is represented in a 4096-dimensional space, which we visualize in **Figure 1A** along the first three principal components for a sample of 500 images. Within this multi-dimensional space, some images are nearby—eliciting similar activation profiles across the set of deep neural network units, while other images are farther apart— eliciting more distinct activation profiles. The set of all natural images in this space comprise the data manifold.

**Figure 1.**
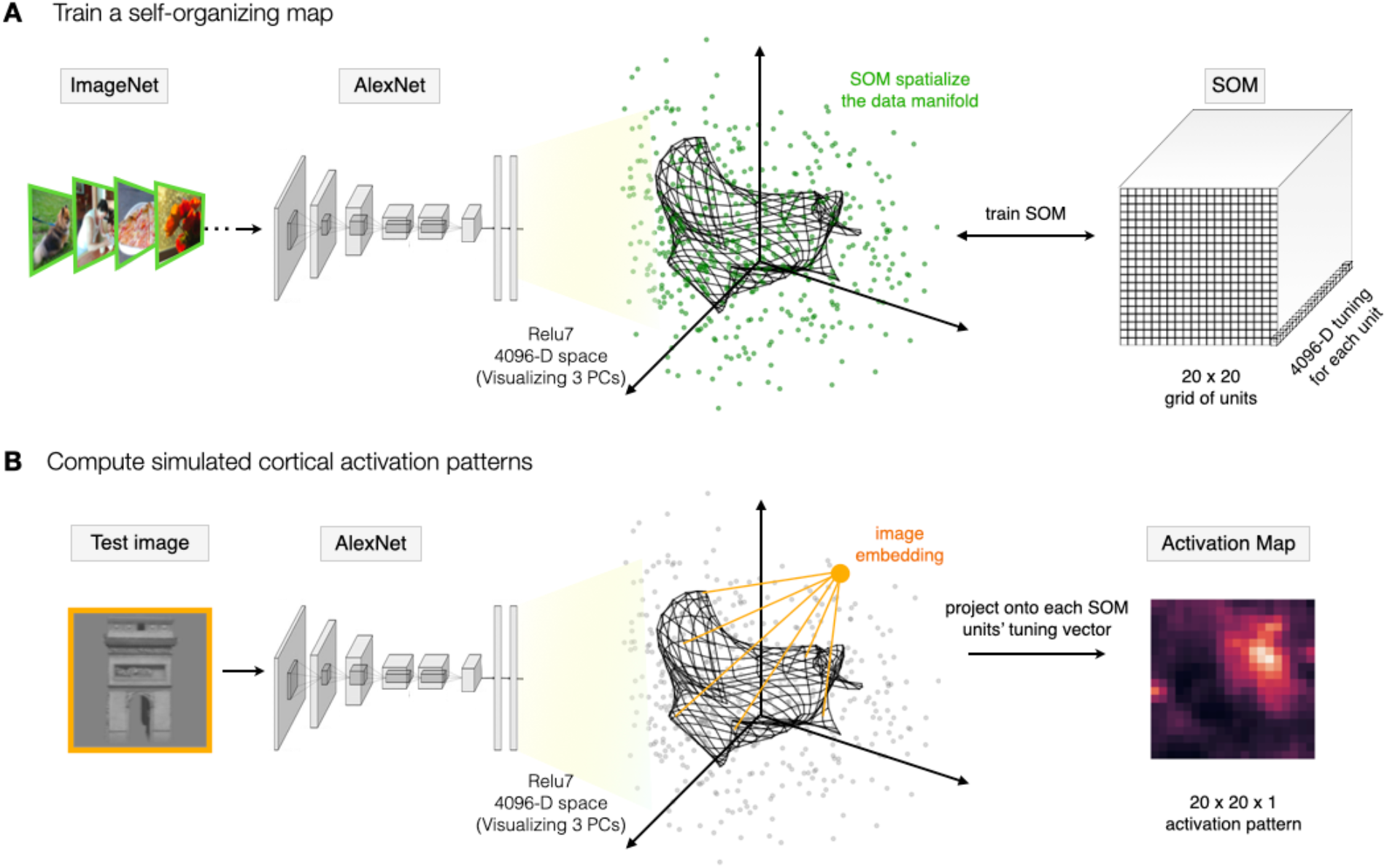
(A) A self-organizing map (SOM) is appended to a pre-trained Alexnet, following the relu7 stage. The relu7 layer is a 4096-dimensional feature space, visualized here along the first three principal components, where the green dots reflect the embedding of a sample of ImageNet validation images. The final SOM layer consists of a two-dimensional map of units of size 20×20, each with 4096-dimensional tuning (depicted as a black grid). During training, the tuning curves of these map units are updated to capture the data manifold of the input images (i.e. the set of green dots). (B) To compute the spatial activation map for any test image, the image is run through the model and the relu7 embedding is computed. Then, for each map unit, the projection of the image embedding onto the tuning vector is computed (conceiving of these tuning vectors as carrying out a filter operation), and this value is taken as the activation of this map unit to this image.

Next, we add a self-organizing map (SOM) layer, which can be conceived of as an additional fully connected layer, where the tuning of each unit of the SOM is a weighted combination of the relu7 features. These tuning vectors of SOM units are trained with the goal of smoothly capturing the data manifold. Specifically, the algorithm projects a 2-dimensional grid of units into the relu7 space, learning tuning curves for each unit such that unit with nearby tuning in the relu7 representational space are also spatially nearby in the grid of map units. Further, the algorithm is designed to ensure that the collective set of map units have close coverage over the entire data manifold. Thus, if there are parts of this feature space that are occupied by natural images, there will be some map units tuned near that part of the representational space. And, if there are combinations of relu7 feature activations that no natural images ever activate, then no SOM units will have tuning curves that point to that part of the representational space. In this way, the SOM transforms the implicit representation of natural images embedded in the feature space to be an explicit map of the data manifold.

The SOM was trained with an iterative algorithm, following standard algorithm procedures (Kohonen, 1990; see **Methods** for details). Note that the specifics of the learning algorithm are not intended to be interpreted as a direct mechanistic model of cortical development. To overview, first, the tuning of each SOM unit was initialized in a grid covering the plane of the first two principal dimensions of the relu7 feature space. Next, the tuning of each unit was iteratively and competitively updated to be increasingly closer to the input data samples, while also ensuring that neighboring units in the map are updated towards similar parts of the data manifold. Here the 50,000 images from the validation set of ImageNet (Russakovsky et al., 2015), were run through a pre-trained Alexnet (with no additional deep neural network weight updates), and the activations from the relu7 stage were used as the input data distribution to train the SOM layer. Additional details related to SOM initialization, neighborhood parameters, learning rate, and other parameters guiding the training process are detailed in the **Methods**. At the end of training, the resulting layer is referred to as an SOM or map, which consists of a grid of units (here 20×20), each with a 4096-dimensional tuning curve.

**Figure 1A** provides a graphical intuition, where the tuning of each map unit is projected into the feature space, with SOM map units depicted as a grid of connected points. Here the tuning of the units on the SOM (i.e. their locations in this feature space) are shown at an intermediate stage of training, for clarity. **Supplementary Figure 1** visualizes the SOM at different training stages from initialization to final. **Supplementary Figure 2** plots the quality of the fit of the SOM to the input data as a function of training epochs, as well as the final tuning similarity between all pairs of SOM units as a function of distance on the trained map.

We next established a pipeline to measure a spatial activation profile over the output map, for any given test image (**Figure 1B**). To do so, we pass an image through the pre-trained Alexnet to compute its 4096-d vector in the relu7 space. Then we compute the response of each SOM unit by conceiving of it as a filter, where the activation of each unit is computed based on the tuning-weighted combination of feature activations (see **Methods**). With these procedures in place, we next followed the empirical literature, leveraging the same stimulus sets and analysis techniques used to map the response topography of the ventral visual stream, but here computed over the simulated activations of the SOM. Any emergent tuning and topography of object distinctions are thus present in the implicit similarity structure of the DNN representation.

### Large-scale organization of animacy and real-world size emerges on the simulated cortex

We first tested for the representational distinction between animate vs. inanimate objects. Stimuli from Konkle & Caramazza (2013) were used, which depict animals and inanimate objects in color on isolated backgrounds (120 each, see examples in **Figure 2A**). Response preferences along the ventral surface of the brain show a large-scale organization by animacy—that is, with an extensive swath of cortex with higher activations to depictions of animals (purple), adjacent to an extensive swath of cortex with higher activations to inanimate objects (green; data from Konkle & Caramazza, 2013).

**Figure 2.**
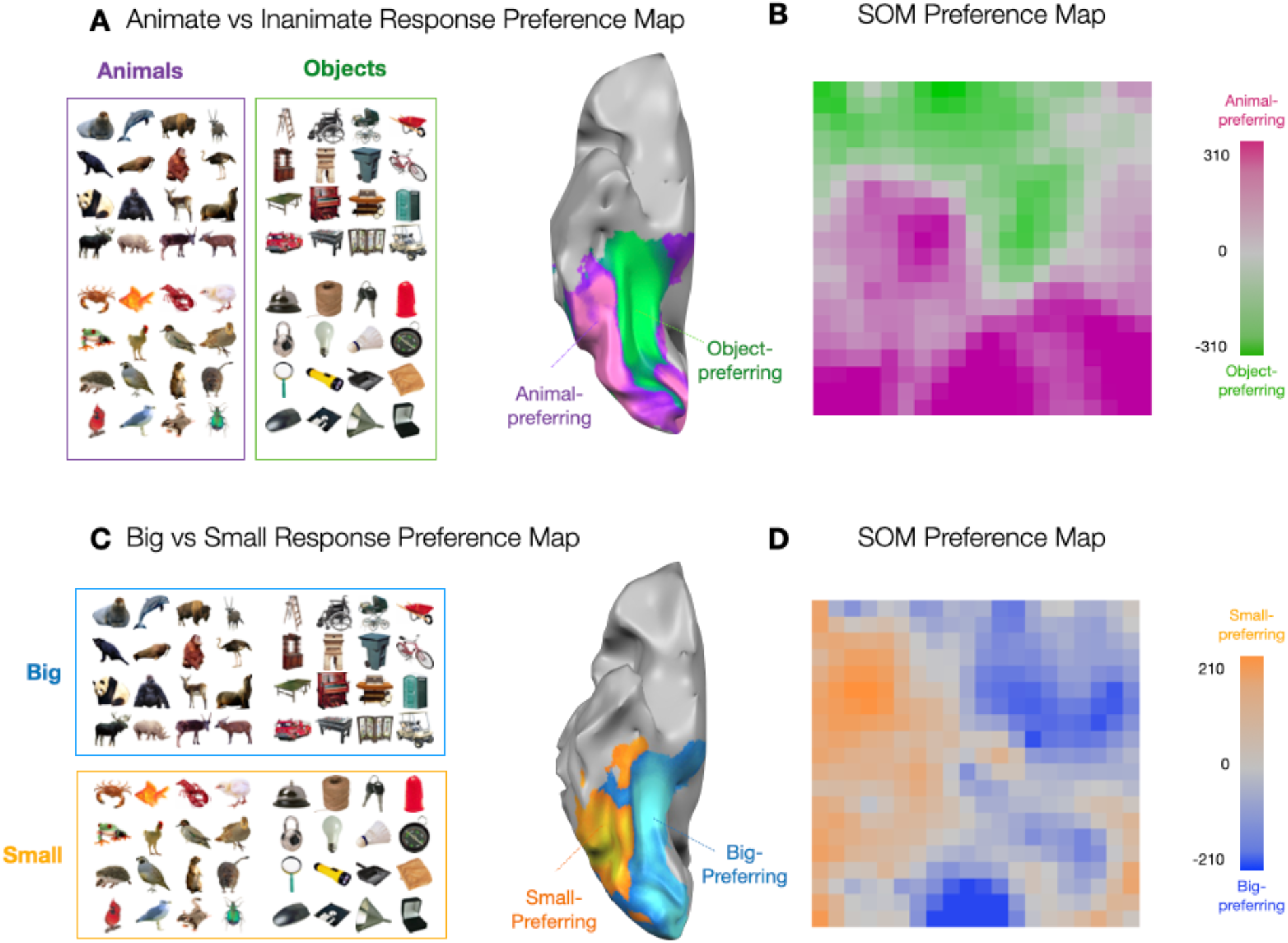
Large-scale organization of animacy and size. (A) Example images of animals and objects shown adjacent to the corresponding brain preference map. Ventral view of a partially inflated hemisphere is shown where regions with stronger responses to depicted objects are shown in green and stronger responses to depicted animals are shown in purple. (B) Each unit of the simulated cortex is colored by its response preference to either animal or object images. (C) Same images as in (A) but now grouped by whether they depict big or small entities in the world adjacent to the corresponding brain preference map. (D) Each unit of the simulated cortex is colored by its response preference for images of big or small entities. Stimuli and brain maps adapted from Konkle & Carramazza, 2013.

For each SOM map unit, we measured the average activation to these same images of animals and objects, and visualized the degree of response preference along the simulated cortical sheet (see **Methods**). The results are shown in **Figure 2B**. Each map unit is colored by whether it has a stronger response to depicted animals or inanimate objects, with stronger response preferences depicted with deeper color saturation. We find that the distinction between animals and inanimate objects reveals many units with preferences for either domain, clustered at a relatively large scale across the entirety of the map. Such an organization was not present when applying the SOM on the same layer’s feature space in an untrained deep neural network, nor in a SOM that was randomly tuned in a 4096-dimensional feature space (**Supplementary Figure 3**).

A second factor that yields large-scale topographic distinctions along the cortical surface of the human ventral visual stream is that of real-world size, shown in **Figure 2C** (Konkle & Oliva, 2011; Konkle & Caramazza, 2013; Julian et al., 2017). That is, there is an extensive swath of cortex that responds more to depicted entities that are typically big in the world (e.g. chairs, tables, landmarks, body-sized or bigger) and an adjacent swath of cortex that responds more to depicted entities that are typically small in the world (e.g. shoes, mugs, tools, and other hand-held manipulable objects), even when these images are presented to the observer at the same visual size.

To visualize the topography of real-world size preferences across the SOM, the same stimuli from Konkle & Caramazza (2013) were used, but instead grouped by size. The size preference map of the SOM again shows a relatively large-scale organization of this factor, with map units showing stronger activations to either big or small entities, clustered at a relatively large scale across the entirety of the map. Such a large-scale organization of response preferences was not present when applying the SOM on the same layer’s feature space in an untrained deep neural network, nor in a SOM that was randomly tuned in a 4096-dimensional feature space (**Supplementary Figure 3**).

These analyses reveal that the distinctions between depicted animate and inanimate objects, and between big and small entities, are related to the major factors of the feature space learned in the deep neural network. For example, it could have been the case that units with animal and object response preferences were tightly interdigitated, or that there were many map units with relatively weak response preferences and only a few with strong domain preferences. Previous empirical work has clearly demonstrated that the animate/inanimate distinction is known to be a major factor in the geometry of both human and non-human primate representation along the ventral stream (Kriegeskorte et al., 2008); here the self-organizing map reveals this property of the deep neural network representational structure in a spatialized format, as a large-scale organization of the response landscape.

### Mid-level visual feature differences underlie the animacy and size organizations

Even though different regions of the brain are systematically activated by images of animals or objects of either big or small sizes, this result does not therefore directly imply that these map units are driven by something very abstract about what it means to be animate or inanimate, big or small. Rather, increasing empirical evidence indicates that responses along this purportedly “high-level” visual cortex have a significant degree of tuning at a more primitive visuo-statistical level (e.g. Long et al., 2018, Jagadeesh & Gardner, 2022; Coggan et al., 2016; Donald & Bonner, 2020). To this end, the next signature of ventral stream topography that we probed is its sensitivity to images with more primitive “mid-level” image statistics preserved (Long et al., 2018).

Long et al., 2018 created images using a texture synthesis algorithm (Freeman & Simoncelli, 2011), which preserved local texture and coarse form information of the original animal and object images, but which were sufficiently distorted to be empirically unrecognizable at the basic level (e.g. lacking clear contours, three-dimensional shape; example stimuli are shown in **Figure 3A**).

**Figure 3.**
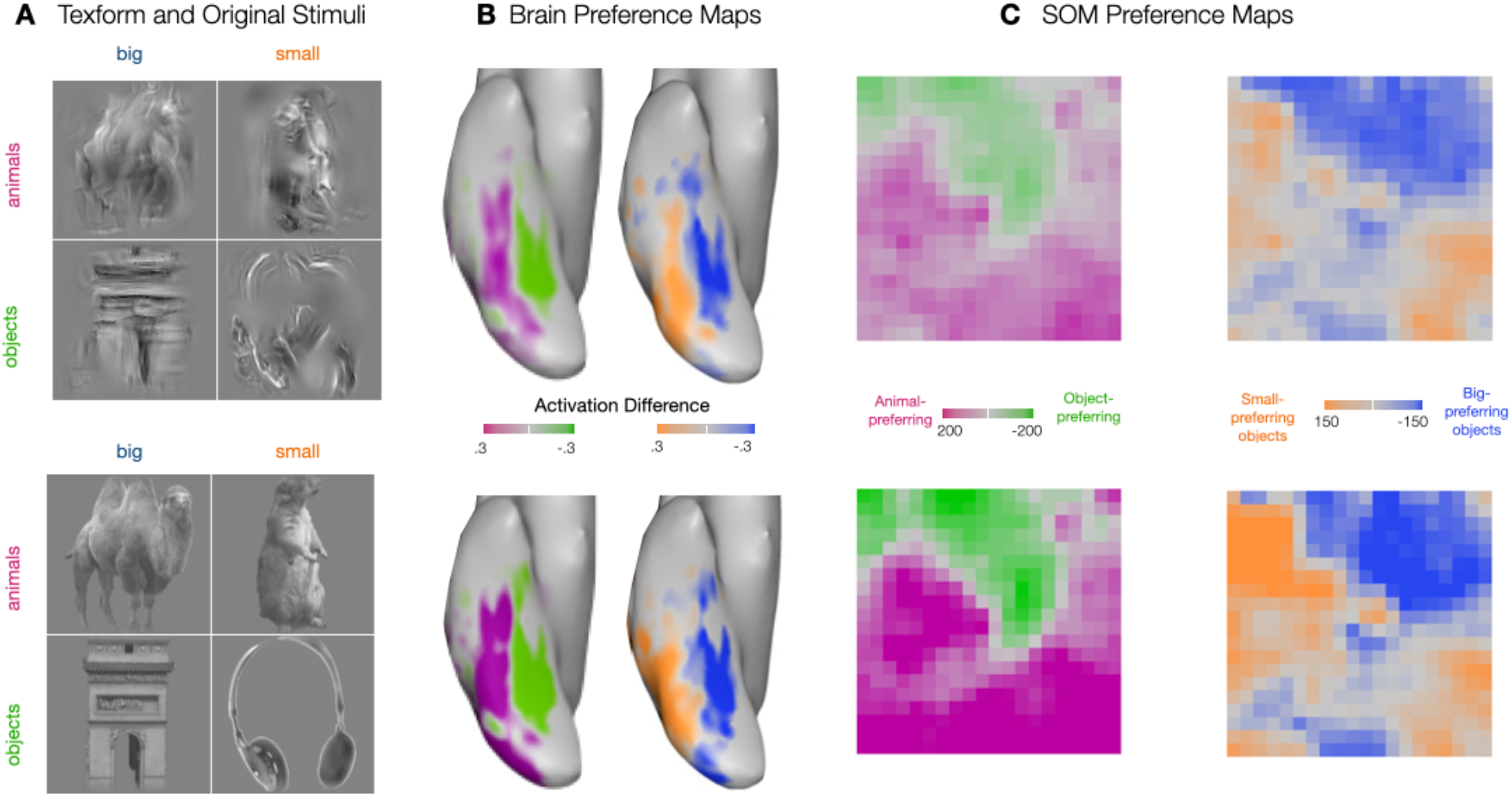
(A) Texform images (upper) generated using a texture synthesis algorithm from recognizable images (below) of 30 big objects, 30 small objects, 30 big animals, and 30 small animals. (B) Preference maps for animacy and size for stimuli shown in (A) along the occipitotemporal cortex. The limits of the color bar reach full saturation at an absolute value of 0.3 reflecting the beta difference computed from an individual’s GLM. (C) Preference maps for animacy and size on the simulated cortex, for texform and original stimuli. Each unit of the simulated cortex is colored based on their preference for animacy, i.e.animals vs objects and for size, i.e. big vs small objects (purple for animals and green for objects in the animacy map and orange for small objects and blue for big objects). Stimuli and brain maps are adapted from Long et al. (2018).

However, these “texform” images still evoked systematic responses along even the later stages of the ventral visual stream. Further, cortex with a preference to animate vs. inanimate recognizable stimuli showed the same large-scale organization in response to texforms, as shown in **Figure 3B**. The same held for real-world size.

To test for these signatures in the SOM, we used the same stimulus set as in the neuroimaging experiment, which consisted of 240 gray-scaled, luminance-matched images (120 originals and 120 texforms, each with 30 exemplars from big, small, animate and inanimate objects). **Figure 3C** shows the corresponding preference maps for texform images and original images, for both animals vs. objects and big object vs. small object contrasts. We find that the mid-level image statistics preserved in texforms are sufficient to drive a near identical large-scale organizations across the SOM (correlation between original and texform maps: animacy r=0.93, p< 10^−5^; size r=0.85, p< 10^−5^).

Thus, these results provide further corroborative evidence that it is possible to have a large-scale organization that distinguishes animals from objects and big objects from small objects without requiring highly abstract (non-visual) features to represent these properties. Instead, this seemingly high-level organization can emerge from visuo-statistical differences learned by deep neural networks, that are particularly reliant on coarsely localized textural features.

### Category-selectivity for faces and scenes

Seminal early findings of ventral visual stream organization also discovered and mapped a small set of localized regions of cortex that have particularly strong responses for some categories of stimuli relative to others, e.g. for faces, scenes/landmarks, bodies, and letter strings (e.g. see **Figure 4B**, Kanwisher et al., 1997; Puce et al., 1996; McCarthy et al., 1997; Epstein & Kanwisher, 1998; Aguirre et al., 1998; Downing et al., 2001; Peelen & Downing, 2005; McCandliss et al., 2003; Polk et al., 2002). Some theoretical accounts of these regions consider these as independent and unrelated functional modules, implicitly assuming no direct relationship between them (Kanwisher, 2010; Zeki, 1978). However, the integrated feature space of the deep neural network allows us to consider an alternate hypothesis that face- and scene-selectivity might naturally emerge as different parts of a common encoding space—one whose features are designed to discriminate among all kinds of objects more generally (Konkle & Caramazza, 2013; Bao et al., 2020; Vinken et al., 2022; Prince & Konkle, 2020; Khosla & Wehbe, 2022). If this is the case, these categories would drive responses in a localized part of the feature space, which would emerge as a localized cluster of selective responses in the SOM.

**Figure 4.**
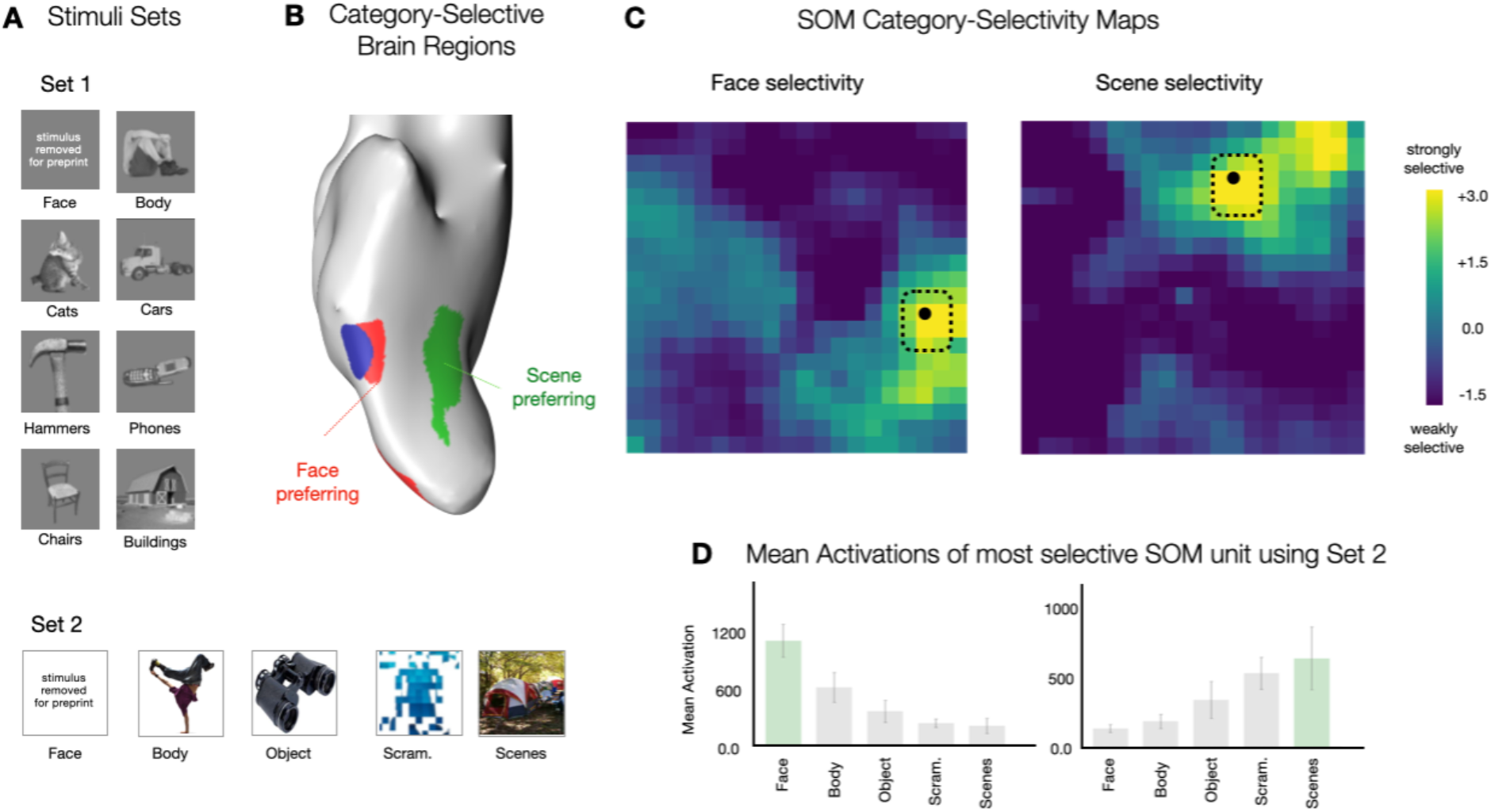
(A) Example images from two Stimuli sets: Stimuli Set 1 containing luminance matched grayscale images from 8 different categories – faces, bodies, cats, cars, hammers, phones, chairs, and buildings; Stimuli Set 2 containing colored images from 5 different categories – faces, bodies, objects, scenes, and scrambled images – on a white background. (B) Ventral view of inflated cortical surface of one individual, highlighting with a face-selective region in red, a body-selective region in blue, and scene-selective region in green. (C) Face-selectivity and Scene-selectivity maps are shown, reflecting a d-prime measure computed over responses to images from Stimuli Set 1. Each unit is colored based on its selectivity for the target category vs. the remaining categories. (D) The most face- and scene-selective map unit was identified, and responses were measured for independent images from Stimulus Set 2. Bar plots show the mean activations of the face-selective map unit (left) and scene-selective map unit (right). Stimulus sets were from Cohen et al., 2017 and Konkle & Caramazza, 2013; and brain maps are adapted from Konkle & Caramazza, 2013.

To explore this possibility, for each map unit, we measured its mean response to images from two different localizer sets (Stimulus Set 1: gray-scaled luminance-matched images of faces, bodies, cats, cars, hammers, phones, chairs, and buildings images; 30 images per category; see example images in **Figure 4A;** Cohen et al., 2017; Stimulus Set 2: 400 color images of isolated faces, bodies, objects, scenes, and scrambled objects on a white background, 80 images per category; Konkle & Caramazza, 2013; see example images in **Figure 4A**). Next, for each unit we calculated the selectivity magnitude, a measure of the d-prime score reflecting the difference between, for example, the response magnitude for all face images, compared with the response magnitude for all non-face images from the set (see **Methods**).

**Figure 4C** plots the selectivity maps for both face and scene selectivity measures, computed over Stimulus Set 1. We find that there are map units with relatively strong selectivity to faces and scenes, clustered in different parts of the SOM. These units showed strong categorical separability (e.g. all face images within the image set were the strongest activating images for the most face-selective unit, while all building images were the strongest activating images for the most scene-selective unit). As a further test of generalizability, we measured the response of the most face- and scene-selective unit in the map to an independent stimulus set, which has different image characteristics. These units again show the strongest response to their preferred category (**Figure 4D**). Finally, the same results were obtained with an alternative *selectivity-index* metric for computing category-selectivity (**Supplementary Figure 4**).

These analyses demonstrate that face and scene regions can naturally emerge in a smoothly mapped DNN feature space, one whose features are learned in service of discriminating many kinds of objects. Thus, these results provide computational evidence for a plausible alternative to the theoretical position that distinct, domain-specialized mechanisms are required for specialized regions with category selectivity to emerge.

### Macro- and Meso-Scale Organization

In the human brain, there is a systematic relationship between the locations of the meso-scale category-selective regions and the response preferences of the surrounding cortex (Konkle & Caramazza, 2013; Weiner & Grill-Spector, 2014). Specifically, the face-selective regions fall within and around the larger zones of cortex that have a relatively higher preferential response to depicted animals, while scene-selective regions fall within zones of cortex that have a relatively higher preferential response to depicted inanimate objects. In the simulated cortex, we find that the same topographic relationship naturally emerges.

**Figure 5A** shows the SOM animate vs inanimate preference map, alongside maps of face- and scene-selectivity, computed for the two different stimuli sets. Qualitative inspection reveals that units with the strongest face-selectivity are located within the region of the map with animate-preferring units and units with the strongest scene-selectivity are located within the region of the map with inanimate-preferring units.

**Figure 5.**
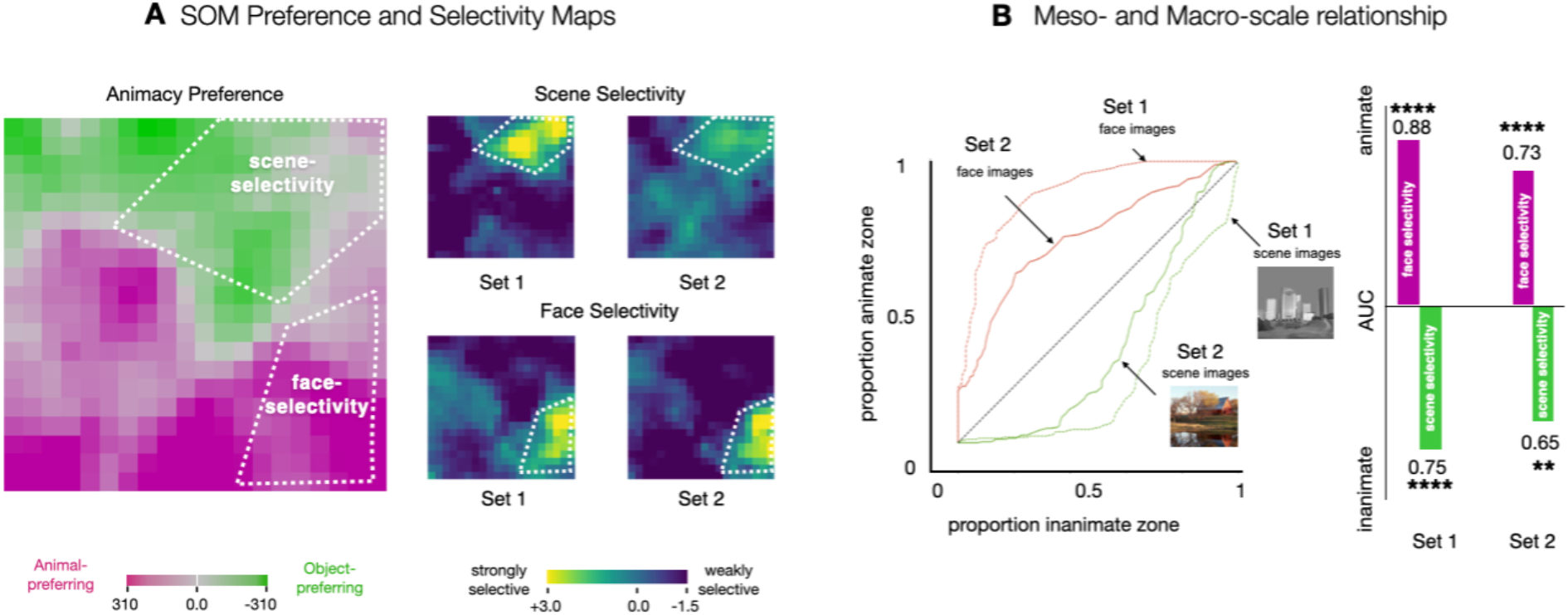
(A) Preference and Selectivity maps. White lines visually demonstrate where the most face-and scene-selective regions fall in reference to the animacy preference zones on the simulated cortex (B) AUC analysis to quantify the category-selective overlap with the preference zones. On the left, we see ROC curves for faces and scenes for both the stimuli sets. These curves reflect how each of the preference zones fill up as increasing number of map units on the SOM get included, starting from the most selective. On the right, we compute area under the curve for the ROC curves.

To quantify the relationship between category-selective maps and the animate-inanimate preference maps, there is a challenge of what threshold to pick to define a ‘category-selective’ region, in order to compute its degree of overlap with the animate-preferring and inanimate-preferring units. To circumvent this issue, we used a receiver operating characteristic (ROC) analysis, following the procedures used in Konkle & Caramazza, 2013 (see **Methods**). This method sweeps through all thresholds, and quantifies where the most selective face units are located, as a proportion of whether they fall in the animate or inanimate zones. By varying the selectivity cut-off threshold (from strict to lenient), this method traces out an ROC curve between (0,0) and (1,1), where the area between this curve and the diagonal reflects how strongly the most selective map units falls within one zone (or the other). Specifically, **Figure 5B** plots the ROC curves and area-under-the-curve (AUC) measures. The face-selective units mainly fall in the animate zones (Set 1: Animate AUC=0.88, p< 10^−5^10^−5^; Set 2: Animate AUC=0.73, p< 10^−5^) while the scene-selective units within the inanimate preferring zone (Set 1: Inanimate AUC=0.75, p<10^−4^; Set 2: Inanimate AUC=0.65, p<10^−2^).

These analyses over the SOM recapitulate previous findings in the brain, highlighting the systematic situation of category selective units within the context of the large-scale organization. As such, they provide computational plausibility for the theoretical position that in the human brain, category-selective regions are not independent islands, but instead, are meaningfully related to each other and to the less-selective cortex just outside them, as part of a unified representational space.

### Divergence between brain and model response topographies

While we have emphasized the topographic signatures that converge between the organization of human object-responsive cortex and the SOM of the penultimate AlexNet layer, there are also clear cases of divergence, at both macro- and meso-scales. Specifically, these differences are evident when considering (1) the interaction between animacy and real-world size properties, and (2) considering which categories show more localized vs distributed selectivity.

The first major difference is related to the way the feature tuning of the DNNs span the animacy and object size distinctions, compared to the human brain. In the simulated cortex, the animacy and object size organizations are relatively orthogonal, e.g. **Figure 2B** shows animate-to-inanimate preferences from the bottom-to-top of the SOM; and small-to-big preference from left-to-right of the SOM. In contrast, as can be seen in the brain organizations in **Figure 2A**, both the inanimate-to-animate and big-to-small contrasts actually evoke a very similar spatial organization along the ventral visual stream, with preferences that both vary from medial-to-lateral.

Konkle & Caramazza (2013) delineated how these two organizations fit together in the human brain, revealing a “tripartite” organization of object tuning (**Figure 6A**). Specifically, they observed that there are three parallel zones of cortex with stronger responses for either depicted big objects, all animals (independent of size), and small objects. Put another way, big and small animals activated relatively similar large-scale patterns across the cortex. The SOM, in contrast, shows an organization with clearer 4-way separability among these conditions (**Figure 6A**). That is, there are zones of SOM map units with a relatively stronger response to either small objects, big objects, small animals, or big animals. This lack of tripartite structure is also evident in the representational geometry of the deep net **(Supplementary Figure 7A)**, highlighting that this divergence is not an artifact of the self-organization process but is inherently present in the structure of the deep net feature space itself.

**Figure 6.**
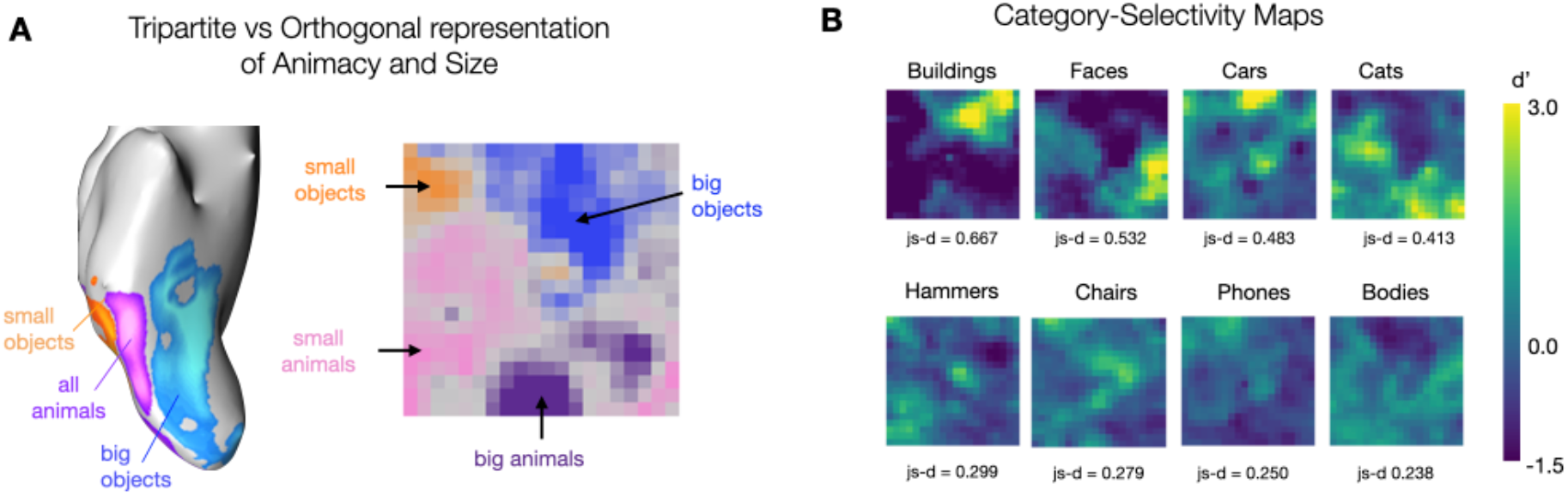
(A) Left: 3-way preference map in the occipitotemporal cortex among big objects, all animals, and small objects (adapted from Konkle & Caramazza, 2013). Right: 4-way preference map in the SOM among big objects, small objects, big animals, and small animals using the same stimuli as used in Konkle & Caramazza, 2013. (B) Category-selectivity maps computed using the d’ measures with each of the 8 categories from stimulus set used in Cohen et al., 2017 serving as the target, and the other 7 categories serving as the non-target images. These plots are organized by an approximate estimate of their non-uniformity, calculated with a js-distance score reflecting on how strongly the distribution of selectivity scores deviates from a distribution of uniform selectivity.

The second divergence between the cortical topography and the SOM of the dnn feature space is related to category-selective signatures across different categories. In the human brain, no highly selective and circumscribed regions that have been mapped for cars, shoes, or other categories (c.f. Downing et al., 2006). However, in the simulated cortex, there is a different pattern. **Figure 6B** shows selectivity maps for each of the 8 categories in the first stimulus set, computed as the d-prime score between the responses over the target category images, relative to the responses over the non-target category images. Qualitative inspection shows that the SOM does have not strongly localized selectivity for bodies, while it does show localized selectivity for cars (and to some extent cats).

In a subsequent post-hoc analysis, we found that body-selectivity was more evident when excluding faces from the d-prime calculation; doing so reveals units with higher body selectivity located precisely where the face-selective units are (**Supplementary Figures 5**). Further, images of faces and bodies are the maximally activating images for neighboring units on the SOM grid (true across several stimulus sets, see **Supplementary Figures 6C and 6D**), consistent with the anatomical proximity of face- and body-selective regions of the human brain (Weiner & Grill-Spector, 2010; Weiner & Grill-Spector, 2013). Thus, body and face tuning are in similar parts of the feature space, but are less separable in the SOM than is evident in cortical organization.

Taken together, these examples reveal that the dnn feature space, when smoothly mapped, has some of representational-topographic signatures that do not perfectly align with the response structure of the object-selective cortex in the human brain.

### A map of object space

The analyses of the tuning of units on the SOM thus far have focused on activation landscapes to different stimulus conditions, similar to the approach taken in fMRI and other recording methods, which measure and compare brain responses to targeted images. However, the tuning of each map unit in the SOM is specified in a feature space of a deep neural network that is end-to-end differentiable with respect to image inputs. This enables us to leverage computational synthesis techniques to visualize the tuning across the map (Olah et al., 2017). Specifically, for each unit’s tuning vector, we extract derivatives with respect to the image, and iteratively adjust the pixel values (starting from a noise seed image) such that it maximally drives a specific unit of the SOM (see **Methods**).

**Figure 7A** schematizes the self-organizing map, embedded in the high-dimensional feature space of the DNN representational space, and depicted below as a flattened grid of tuned units. For a subset of units systematically sampled across the map (25 units highlighted in black), **Figure 7B** shows the corresponding synthesized image that maximally drives these units. **Supplementary Figure 8** shows the synthesized images for all the map units on the SOM. At a glance, these images seem to capture rich textural features, consistent with both what is now known about the nature of the feature representations in DNNs (Geirhos et al., 2018; Hermann et al., 2020). A more detailed inspection shows that the nature of the image statistics captured across the map vary systematically and smoothly, e.g. with synthesized maximally-activating images that clearly are more animal-like or more scene-like in different parts of the map. As a complementary visualization, in **Supplementary Figure 6**, we show the image that maximally drives each map unit, computed over different stimulus sets, including those from Bao et al., 2020.

**Figure 7.**
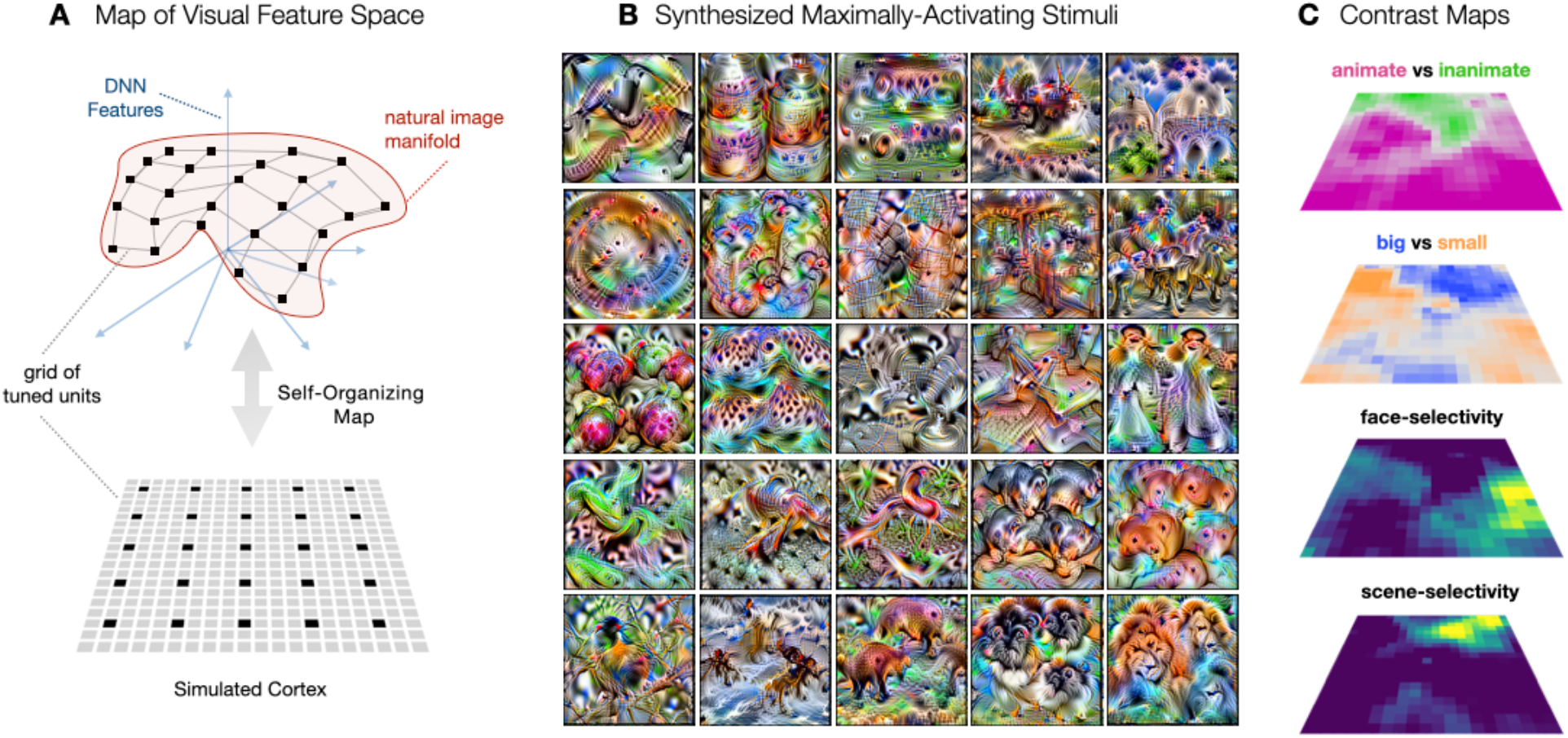
(A) Manifold of natural images formed by the images’ dnn features shown in red. The axis demonstrates each of the 4096 dimensions in the relu7 activations of the dnn. The gray connected lines depict the simulated cortex trying to hug this image manifold and the black points depict units on the map/simulated cortex. This simulated cortex can be described via a 2d grid of connected units, i.e. the SOM (B) Images synthesized using gradient ascent to maximally activate the map units highlighted in (A) via black points(C) Entire spatial hierarchy learned by the simulated cortex – animacy and size preference and face- and place-selectivity

**Figure 7C** provides further context for understanding the map of object space, showing how the organizations of animate vs. inanimate, big vs. small entities, face-selectivity, and scene-selectivity, all are evoked from the same spatialized feature space. This visualization further helps clarify how these preferences for animate vs. inanimate objects, big vs. small entities, and localized regions for faces and scenes, can be related purely to different image-statistics (as any more abstract, non-visual level of representation is beyond the scope of this deep neural network).

Finally, we conducted several SOM variations to examine the robustness of these representational-topographic motifs. **Supplementary Figure 9** shows little to no effect of changing or increasing the number of images used to initialize the SOM tuning. **Supplementary Figure 10** shows that SOMs with approximately 2-3 times more units also showed the same motifs.

## DISCUSSION

Here we used a self-organizing map algorithm to spatialize the representational structure learned within the feature space of a deep neural network trained to do object categorization. This method yields a two-dimensional grid of units with image-computable tuning, that reflects a smooth mapping of the data-manifold in the representational space. We tested whether several hallmark topographic motifs of the human object-responsive cortex were evident in the map, finding several convergences. First, large-scale divisions by animacy and real-world object size naturally emerged. Second, the same topographic organizations were elicited from unrecognizable “texform” images, indicating the feature tuning is sensitive to mid-level visual statistical distinctions in these images. Finally, clustered selectivity for faces and scenes naturally emerged, without any specialized pressures to do so, and was situated systematically within the broader animacy organization, as in the human brain. However, the simulated cortex did not capture all macro- and meso-scale signatures. For example, it contained an orthogonal rather than a tripartite representation of animacy and size, and lacked localized body-selective regions, leaving open questions for what is needed to learn an even more brain-like organization. Theoretically, this work provides computational plausibility towards a unified account of visual object representation along the ventral visual stream.

### Implications for the biological visual system

After two decades of functional neuroimaging research charting the spatial structure of object responses along the ventral visual stream, it is clear that there is a stable, large-scale topographic structure evident across people; however, the guiding pressures that lead to this stable organization are highly debated (Op de Beeck et al., 2008; Arcaro & Livingstone, 2017; Arcaro et al., 2019; Arcaro et al., 2009; Dahaene & Cohen, 2007; Saygin et al., 2016; Peelen et al., 2014; Peelen et al., 2013; Hasson et al, 2003; Mahon & Caramazza, 2011). On one extreme, for example, the nature of the tuning and the locations of category-selective regions are primarily driven by specialized pressures that are innate and non-visual in nature, with supporting evidence from distinct long-range connections beyond the visual system, and co-localized functional activations in the congenitally blind (Saygin et al., 2012; Saygin et al., 2016; Osher et al., 2016; Mahon & Caramazza, 2011; Peelen et al., 2014; Peelen et al., 2013; Striem-Amit & Amedi, 2014; Striem-Amit et al., 2012; Konkle & Caramazza, 2017). On the other extreme, it is the experienced statistics of the visual input, scaffolded from an initial retinotopic organization and generic learning mechanisms, that are primary drivers of the organization in the object-selective cortex (Arcaro & Livingstone, 2017; Arcaro et al., 2019;Arcaro et al., 2017; Dahaene & Cohen, 2007; Dahaene et al., 2015; Konkle & Oliva, 2012; Konkle & Caramazza, 2013; Konkle & Alvarez, 2022; Plaut & Behrmann, 2011). What can the present modeling work contribute to this debate?

Here we suggest that, by probing the representational signatures evident in this model, we gain traction into what kind of object distinctions can emerge from the experienced input, without requiring category-specialized pressures. That is, the network is capable of extracting the regularities in input distributions, reformatting them into a code that can support downstream behavior like object categorization. For example, the Alexnet architecture we used does not have any explicit learning mechanisms devoted for some special categories (e.g. branching architectures that are trained only with faces; Dobs et al., 2022). Similarly, the SOM also does not have any category-specific learning rules. In this way, our model leverages a relatively generic set of inductive biases that guide the structure of the learned visual feature space. In this way, rather than thinking of this deep neural network as an exact model of the visual system, we can think of it instead as a functionally powerful representation learner.

On this framing, the fact that the SOM shows a large-scale organization by animacy and object size, without explicit connectivity-driven pressures or domain-specific learning mechanisms that enforce these groupings, means that these “high-level” distinctions can emerge directly from image-statistical differences in the input. The results with texforms corroborate this interpretation. Further, we show that even clustered face-selectivity and scene-selectivity emerge—indicating that depicted faces and scenes have a particularly focal and separable location in the deep neural network feature space—and need not be attributed to specialized learning pressures. Certainly, this result does not provide direct mechanistic evidence for the experience-based formation of these regions in the brain. But, while experience-based accounts formerly could only speculate that certain object category distinctions could emerge from input statistics alone, this work now provides clear support for the sufficiency of image statistics to form a basis towards the emergence of these distinctions.

It is an open question whether the divergent topographic motifs we observed are a result of critical missing architectural or algorithmic components that help guide visual system topography, or whether these can be attributed to differences in the experienced input statistics between models and humans. Indeed, the more face images in the input set, the denser that part of the data manifold will be, and the more SOM-territory will be devoted to that part of the feature space. We hypothesize that different input statistics will ultimately be able to account for these differences, perhaps coupled with different downstream behavioral goals (e.g. self-supervised objectives), without requiring body-specific learning mechanisms to guide the development of body-selective regions (and anti-car mechanisms to reduce the development of car-localized regions). To this end, we explored the organization of an Alexnet trained on the Ecoset database (Mehrer et al., 2021), which has a different distribution of categories—in this model, we did observe more of a tripartite structure, but still did not find localized body-selective regions (**Supplementary Figure 7B**). In this way, this work also introduces a possible new method to visualize the impact of different input datasets, architectures, and tasks in shaping the format of the learned representation.

Finally, it is important to acknowledge that there are also many other empirical signatures of object topography, which these models are not yet directly equipped to test. For example, object topography along the cortex in humans is “mirrored,” with duplicated selectivity on the ventral and lateral surface (Taylor and Downing, 2011; Hasson et al., 2003; Kravitz et al., 2013). This duplication has been hypothesized to emerge from extensions of adjacent retinotopy, reflecting the divisions of the upper and lower visual field, (though the influence of non-duplicated area MT on the lateral surface has also been hypothesized). More generally, there is an extensive trove of empirical and anatomical data, coupled with existing hypotheses about their role in driving the tuning and topography along the ventral visual stream, simply awaiting the advancing frontier of image-computable modeling frameworks to explore these theories. Until then, we offer that by considering this deep neural network model and SOM as a representational system, rather than a direct model of the visual system, still allows for computational insights into the possible pressures guiding the organization of the ventral visual stream.

### Modeling Cortical Topography

How does the approach taken here relate to concurrently developed techniques bringing spatialized responses to deep neural networks (Lee et al., 2020; Blauch et al, 2022; Keller et al., 2021; Zhang et al., 2021)? Across the set of approaches, all seem to be conceiving of the problem at different levels of abstraction, and test for different signatures. For example, Lee et al., 2020 conceive of the early convolutional layers as already having topographic constraints, while the fully-connected layers do not; they arranged the fully-connected units in a grid and added a spatial correlation loss over the tuning during model training, in addition to the object categorization objective. They found clusters of face-selective units that were connected across the fully-connected layers—they did not however probe for animacy, object size, or other category-selective regions. Blauch et al., 2022 instead dropped the fully connected layers, and instead added three locally connected spatialized layers, with coupled excitatory and inhibitory processes. When trained on faces, objects, and scenes, these layers show increasing clustering to these categories. In both approaches, topographic constraints are directly integrated into the feature learning process.

In contrast, we cast the problem of topography as one of data-manifold mapping, which is more closely related to the approaches taken by Keller et al., 2021 and Zhang et al., 2021. Keller et al., 2021 trained a topographic variational autoencoder which, like our SOM, was also trained on from the features from of a pre-trained Alexnet model (though appended after the final convolutional stage). This topographic layer is also a grid of units (though, with a circular topology), initialized into the deep net feature space, and trained to maximize the data likelihood using an algorithm related to independent component analysis. Similarly, Zhang et al., 2021 also leveraged a pre-trained Alexnet (though, they used the final output layer, first reducing it to 4 dimensions using PCA), and then trained an SOM where each unit has a 4-dimensional tuning curve. Both these approaches probe the resulting tuned map with some of the same stimulus sets as in the present work, though we all used different analysis methods to quantify the spatial organization, resulting in some differences (e.g. both Keller et al., 2021 and Zhang et al., 2021 report the presence of body-selective regions). As a whole, these methods use a topographic layer to reveal the untangled data manifold of a pre-trained feature space, rather than to constrain the learning of the features themselves.

Given this formulation of topography, we do not take the present model as a mechanistic model of cortical topographic development. To this end, we see the relevant level of abstraction to approach the mechanistic model as one that takes on the full topographic challenge, learning the growth rules to connect a grid of units into a useful hierarchical network architecture (likely leaning on an eccentricity-based scaffold and the activity of retinal waves to initialize the architecture). However, many other approaches are also possible which reflect different abstractions, e.g. incorporating differentiable SOM stages after each hierarchical layer block, to allow for an interplay between feature learning and data-manifold mapping during representation formation.

Finally, complementing these computational approaches, there is a clear need to develop quantitative metrics for comparing topographic activation similarity, which take into account distance on a cortical sheet (e.g. Wasserstein distance). Recent open, large-scale condition-rich fMRI datasets are now available (e.g. NSD dataset, Allen et al., 2022; THINGS dataset, Hebart et al, 2019, 2022) which can enable the development of cortical topographic metrics beyond these macro- and meso-scale signatures probed for here. Thus, going forward, there is clear work to do towards mapping these computational models more directly to the cortex (c.f. Zhang et al., 2021), and assessing how they succeed and fail at capturing the systematic response structure to thousands of natural images across the cortical surface.

## METHODS

### Spatializing the representational space of a deep net with a self-organizing map

#### Input Data and SOM Parameters

We applied a Kohonen Self-Organizing Map algorithm (Kohonen, 1990) to the multi-dimensional feature space of the relu7 stage of a pre-trained AlexNet (Krizhevsky et al., 2012) sourced from the Torchvision (PyTorch) model zoo (Paszke et al., 2019). The input data is a set of *p* points encoded along *f* feature dimensions. Here, the *p* points reflect the 50,000 images from the ImageNet validation set, and the *f* dimensions reflect the 4096 features from the relu7 stage of the network i.e. *f* ∈ {*f*_1_, *f*_2_, …, *f*_4096_}. Additionally, we specify the number of SOM units (here 400 units) as an input parameter, and set additional training hyperparameters related to the number of training epochs, and how the learning rate and map neighborhood influence changes over the course of map training, detailed below.

#### SOM Training

The first stage of the algorithm is to define the map shape, and then initialize the tuning for each unit on the map such that the map spans the first two principal components of the input data. Computing the principal components over 50,000 points in the 4096-dimensional input space is computationally intensive; thus, we created a smaller sample of 400 images over which we computed the top two eigenvectors and eigenvalues. In a control analysis, we varied the images and the size of this subset over which the principal components were calculated and found that this choice had negligible impact on the final results (**see Supplementary Figure 9)**.

The first step is to determine the aspect ratio of the SOM, based on the ratio of the top two eigenvalues. In the case of the *relu7* feature space, the aspect ratio of the data was ∼ 1, thus the input parameter of **400 map units** lead to the construction of a 20 × 20 (*W*H*) map grid. Next, each unit in the 20*20 grid is placed in the 4096-dimensional space such that the entire map is centered along the plane formed by the first two eigenvectors, scaled by their respective eigenvalues (see top row of **Supplementary Figure 1**). To scale the eigenvectors, we compute unit vectors along the two principal components and multiply them with the square root of their corresponding eigenvalues. Hereon, we refer to the location of a map unit in the 4096-dimensional space as that unit’s ***tuning*** vector and the set of all map tuning vectors as the ***codebook***, which is of size *W x H x f*, here 20 × 20 × 4096. This method of initialization ensures the map is matched to the relative contributions of the top two major dimensions/axes of variation in the input data and allow for a more consistent embedding in this high-dimensional input space.

After initializing the map tuning vectors, the next stage is to fine-tune and iteratively update these tuning vectors to better capture the input data manifold. All 50,000 images from the ImageNet validation set were used during fine-tuning. The full image set is seen every epoch and the SOM was tuned for a total of **100 epochs**. Within each epoch, the map tuning updates operate over a smaller batch of images. Our batch size was **32 images**. For every image in the batch, we first identify the single SOM unit who’s 4096-dimensional tuning vector is closest to that image’s 4096-dimensional embedding in the deep neural network feature space, using the Euclidean distance metric. This SOM unit becomes the image’s “best matching unit” or BMU (see **Equation 1**).

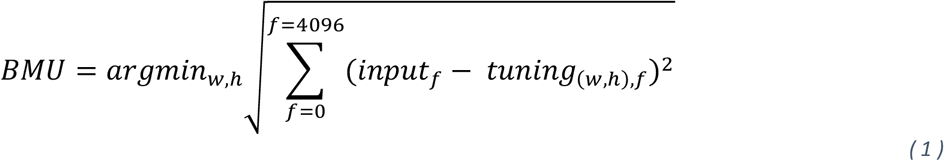

Here ***input***_*f*_ is the image’s dnn activation value on the *f*^*th*^ feature dimension and *tuning*_*(w,h),f*_ is the scalar value, for the *f*^*th*^ feature dimension, on the tuning vector of a map unit that is situated in the *w*^*th*^ row and *h*^*th*^ column of the SOM grid. Hence, the BMU is the SOM unit with the minimum euclidean distance to the image’s feature vector (i.e., ***input***) among all the SOM units. Next, for each of the BMUs (32 per batch), we adjust its tuning vector, and the tuning vectors of other map units that are within a neighborhood of the BMU such that they are closer to the 4096-dimensional location of the corresponding image. This update rule, at a particular time step ***t*** (i.e., *epoch)*, is formulated in **Equation 2**.

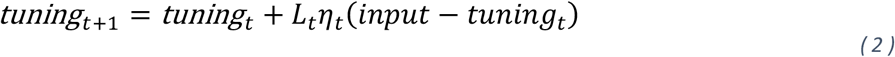

Here the *tuning* vector of each map unit is adjusted towards the *input* based on the learning rate function *L*_*t*_, and the neighborhood function *η*_*t*_. The learning rate (*L*_*t*_) controls the magnitude of the tuning adjustment, which slowly decays to make smaller adjustments over time, following **Equation 3.** The initial learning rate *L*_0_ was set at **0.3** and **T** denotes the total number of epochs (set to 100).

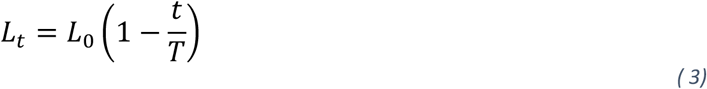

The neighborhood function *η*_*t*_ measures the influence a map unit’s **distance** from the BMU has on that map unit’s learning. Intuitively, units that are closer to the BMU need to be updated more strongly as compared to units that are further away. This is expressed using a gaussian widow (see **Equation 4**) that is centered around the computed BMU with a radius/standard deviation of *σ*_*t*_.

To center the window around the BMU, **Equation 5** is used which computes the L2-distance between a unit present in the *i*^*th*^ row and *j*^*th*^ column and a BMU that situated in the *w*^*th*^ row and *h*^*th*^ column of the SOM grid. It is important to note that this distance is computed directly on the 2d SOM grid and not in the 4096-dimensional input space. This constraint generally encourages neighboring units on the map to encode nearby parts of the high-dimensional input space. For the radius of the neighborhood window, we start with a radius of *σ*_*o*_ that covers approximately half of the map (hence for the map of shape 20*20 it was set at **10**). This radius exponentially decays over the training epochs following **Equation 6**. By starting with a larger neighborhood and gradually shrinking the neighborhood influence, the map is less influenced by image order and batch size, and stabilizes in a smoother larger-scale embedding

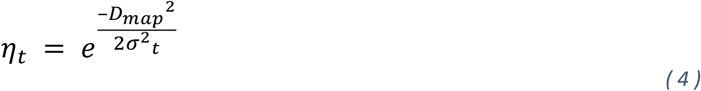

where:

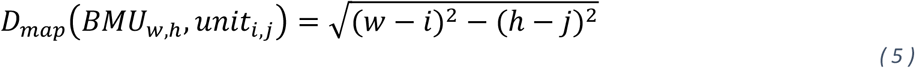

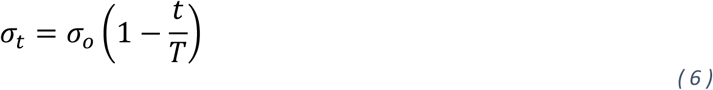

Map tuning updates are made for each batch, with a single epoch completed after all 50,000 images have been presented. At the next epoch (i.e. next time step *t*), the learning rate and neighborhood parameters are updated (using **Equation 5 and 6**) and the process is repeated, continuing for a total of 100 epochs. Due to the decay of the learning rate, the training stabilizes at the end of the total epochs, and we do not find a large differences in the codebook with more training epochs.

A standard measure of map fit to the input data is the Quantization Error (QE), which is the average Euclidean distance between the input image’s dnn features and the tuning of their corresponding computed BMUs. As the map is fine-tuned, this tuning better matches the input data, and the QE decreases. A plot of the QE over epochs is shown in **Supplementary Figure 2A**. In **Supplementary Figure 2B and 2C** we visualize the pairwise tuning of SOM units as a function of their distance on the 2d grid. The tuning similarity reduces as distance on the 2d grid increases as expected via the constraint introduced in **Equation 4**.

At the end of the fine-tuning phase, we have a trained SOM, or “simulated cortex”—a grid of units of shape 20 × 20, each tuned systematically in the high-dimensional space (ℝ^4096^) to encode the data manifold of the input of natural images in the *relu7* feature space of Alexnet.

### Simulated Cortical Activations

To get the activations of new images on the simulated cortex, we pass the image through the pre-trained Alexnet and compute its 4096-dimensional features in the *relu7* space (i.e. *input* vector for that image). Each unit on the SOM also has an associated *tuning* vector in this feature space (ℝ^4096^) and can be conceived of as a filter, i.e. a weighted combination of the dnn features. Thus, we compute the activation of each SOM unit by taking the dot product of that unit’s tuning vector and the image’s relu7 features using **Equation 7**. Across all map units, this creates a spatial activation profile for the image.

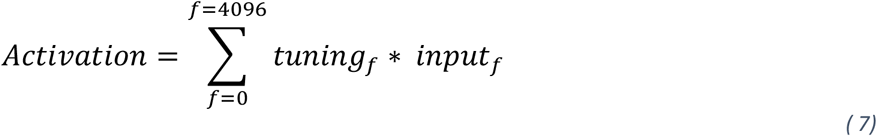

### Stimulus sets

The following stimulus sets were used to probe the spatial topography of the SOM. (1) Konkle & Caramazza, 2013: Animacy x Size images – 240 color images of big animals, small animals, big objects, and small objects (60 each). (2) Long et al., 2018: Original and Texform Animacy x Size images – 120 gray-scaled luminance matched images and 120 corresponding texform images, depicting animals and objects of big and small sizes (30 each) (3) Cohen et al., 2017: Category-Localizer Stimulus Set 1: 240 total gray-scaled luminance-matched images of faces, bodies, cats, cars, hammers, phones, chairs, and buildings (30 each). (4) Konkle & Caramazza, 2013: Category-Localizer Stimulus Set 2: 400 total color images of faces, bodies, scenes, objects, and block-scrambled objects on a white background (80 each).

### Preference maps

Preference maps were created following the same procedures as used in fMRI analysis (e.g. Konkle & Caramazza, 2013). Simulated cortical activations (**Equation 7**) were computed for all individual images from the stimulus set. For each map unit, we computed the average activation for each targeted image condition (e.g. averaging across all animal images or all object images). Next we identify the “preferred” condition, eliciting the highest average activation, and calculated this response preference. For two-way preference maps, the preference strength is the absolute difference between the mean activations of the two categories. For *n*-way contrasts, the preference strength is the absolute difference between the activation of the preferred condition and the second-most activation condition. We visualize the response preferences using custom color maps that interpolate between gray and the target color for each condition, where the color of each unit reflect the preferred category color, and strength of the preference scales the saturation. The mapping between the color palette and the data values are controlled with color limit parameters and were matched across the multiple color maps in the preference map visualization.

### Category-selectivity metrics

To compute maps of category selectivity, we used the following procedure. First, we computed the simulated cortical activations (using **Equation 7**) for all images in the localizer set. Next, for each unit on the map, we computed the mean and variance of activation responses for images from the target category (i.e.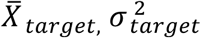) and for all remaining images (i.e. non-target condition; 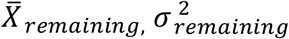)and computed d’ following **Equation 8**.

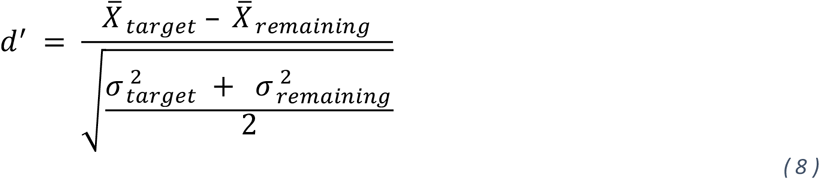

For robustness, we additionally computed another standard measure – Selectivity Index (*SI*) for each map unit, which differs slightly from d’ in how it is normalized (i.e. by the means, rather than the variances), following **Equation 9**. Both metrics yielded convergent results (see **Supplementary Figure 4**).

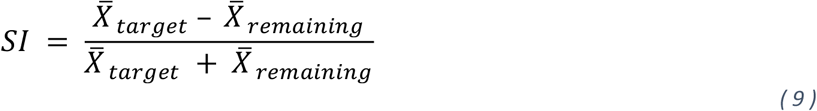

For each map, we also computed a non-uniformity score, based on how different the selectivity map was from a uniform distribution. For each selectivity map, we normalize the d’ scores using a softmax function to get a probability distribution **P** of the selectivity on the map. We then compare this to a completely uniform distribution of selectivity **Q**, using the Jenson Shannon distance following **Equation 10**, where *KL*(*P*||*Q)*is the KL-divergence between distribution P and Q.

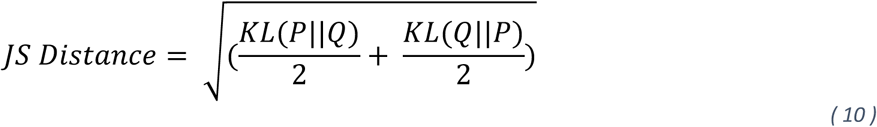

### Comparing selectivity maps and preference maps

To quantify the relationship between category-selective maps and the animate-inanimate preference maps, we used a receiver operating characteristic (ROC) analysis, following the procedures used in Konkle & Caramazza, 2013. The procedure is as follows, described here for the specific case of comparing the map of face-d’ and the map of responses of animate-inanimate preferences. First, the face d’ values are sorted across all 400 map units on the 20*20 grid. For each step in the analysis, the top-most selective units are selected, starting from the top 1% most face-selective, then top 2% most face-selective, and so on, until we consider all 100% of the map units. For each step, we separately compute the proportion of all animate-preferring SOM units and the proportion of all inanimate-preferring units that overlap with these face-selective units. Across all steps of the analysis, an as an increasing number units from the face-selectivity map are considered, the procedure sweeps out an ROC curve between (0,0) and (1,1). For example, if all of the top-most face-selective units were also all animate-preferring units, this curve would rise sharply (indicating rapid filling of the animate-preferring zone), before leveling off. Thus, the area between the curve and the diagonal of this plot (AUC) was used as a threshold-free measure of overlap between face-selectivity and the animacy organization. We computed these ROC curves for both the face- and scene-selective contrasts, computed over both localizer sets.

To measure the significance in this relationship between category-selectivity and the large-scale preference organization we used permutation tests i.e., we iterated through 1000 simulations, and for each simulation, we randomly shuffled the selectivity measure estimates. For each shuffled simulation, we plot the ROC curve across the thresholds and evaluate the AUC measure. The proportion of these simulated AUC’s that are higher than the originally measured (unshuffled) AUC gives us the significance of the measured AUC overlap.

### Representational Geometry and Multi-dimensional Scaling Plots

The 240 images of animals and objects of different sizes was passed through the pre-trained Alexnet and the 4096-dimensional features in the *relu7* space were extracted for each image. Next, pairwise correlations were conducted over these features using the 1-Pearson correlation measure, yielding a representational dissimilarity matrix of 240 × 240 matrix. This matrix was inputted into a standard multi-dimensional scaling (MDS) algorithm with output dimensionality set to 2. Images that are more similarly represented in the dnn feature space are closer to each other in the 2d MDS plot.

### Gradient-based image synthesis

Given that the tuning of each unit on the SOM can be conceived of as a weighted combination of the relu7 features, we can conceptualize an SOM as an additional fully connected layer on top of the relu7 layer with a weight matrix of shape 4096*400 i.e. the 4096-dimensional tuning vector for each of the 400 units on the 20*20 grid of the SOM. This model (i.e. dnn + attached layer from SOM tunings) is end-to-end differentiable with respect to the input images. As a result, we can start with a noise image, and iteratively update it using gradient ascent such that the optimized image increase the output for a selected output unit (which is equivalent to increasing the simulated cortical activation of a unit on the SOM). We use the torch lucent library (https://github.com/greentfrapp/lucent) to synthesize these images.

## ACKNOWLEDGEMENTS

This work was supported by NSF CAREER: BCS-1942438 to T.K. We would like to thank lab members of the Vision Science Lab for their helpful feedback and support during the writing process.

## Notes

### Competing Interest Statement

The authors have declared no competing interest.

### Summary of Updates

First version of the manuscript draft

## REFERENCES

1. Aflalo, T.N. and Graziano, M.S., 2006. Possible origins of the complex topographic organization of motor cortex: reduction of a multidimensional space onto a two-dimensional array. Journal of Neuroscience, 26(23), pp.6288–6297.

2. Aflalo, T.N. and Graziano, M.S., 2011. Organization of the macaque extrastriate visual cortex re-examined using the principle of spatial continuity of function. Journal of neurophysiology, 105(1), pp.305–320.

3. Aguirre, G.K., Zarahn, E. and D’Esposito, M., 1998. An area within human ventral cortex sensitive to “building” stimuli: evidence and implications. Neuron, 21(2), pp.373–383.

4. Allen, E.J., St-Yves, G., Wu, Y., Breedlove, J.L., Prince, J.S., Dowdle, L.T., Nau, M., Caron, B., Pestilli, F., Charest, I. and Hutchinson, J.B., 2022. A massive 7T fMRI dataset to bridge cognitive neuroscience and artificial intelligence. Nature neuroscience, 25(1), pp.116–126.

5. Arcaro, M.J. and Livingstone, M.S., 2017. A hierarchical, retinotopic proto-organization of the primate visual system at birth. Elife, 6.

6. Arcaro, M.J., McMains, S.A., Singer, B.D. and Kastner, S., 2009. Retinotopic organization of human ventral visual cortex. Journal of neuroscience, 29(34), pp.10638–10652.

7. Arcaro, M.J., Schade, P.F. and Livingstone, M.S., 2019. Universal mechanisms and the development of the face network: what you see is what you get. Annual review of vision science, 5, p.341.

8. Arcaro, M.J., Schade, P.F., Vincent, J.L., Ponce, C.R. and Livingstone, M.S., 2017. Seeing faces is necessary for face-domain formation. Nature neuroscience, 20(10), pp.1404–1412.

9. Bao, P., She, L., McGill, M. and Tsao, D.Y., 2020. A map of object space in primate inferotemporal cortex. Nature, 583(7814), pp.103–108.

10. Behrmann, M. and Plaut, D.C., 2013. Distributed circuits, not circumscribed centers, mediate visual recognition. Trends in cognitive sciences, 17(5), pp.210–219.

11. Blauch, N.M., Behrmann, M. and Plaut, D.C., 2022. A connectivity-constrained computational account of topographic organization in primate high-level visual cortex. Proceedings of the National Academy of Sciences, 119(3), p.e2112566119.

12. Chao, L.L., Haxby, J.V. and Martin, A., 1999. Attribute-based neural substrates in temporal cortex for perceiving and knowing about objects. Nature neuroscience, 2(10), pp.913–919.

13. Coggan, D.D., Watson, D.M., Wang, A., Brownbridge, R., Ellis, C., Jones, K., Kilroy, C. and Andrews, T.J., 2022. The representation of shape and texture in category-selective regions of ventral-temporal cortex. European Journal of Neuroscience.

14. Cohen, M.A., Alvarez, G.A., Nakayama, K. and Konkle, T., 2017. Visual search for object categories is predicted by the representational architecture of high-level visual cortex. Journal of neurophysiology, 117(1), pp.388–402.

15. Conway, B.R., 2018. The organization and operation of inferior temporal cortex. Annual review of vision science, 4, p.381.

16. Cowell, R.A. and Cottrell, G.W., 2013. What evidence supports special processing for faces? A cautionary tale for fMRI interpretation. Journal of Cognitive Neuroscience, 25(11), pp.1777–1793.

17. de Beeck, H.P.O., Pillet, I. and Ritchie, J.B., 2019. Factors determining where category-selective areas emerge in visual cortex. Trends in cognitive sciences, 23(9), pp.784–797.

18. Dehaene, S. and Cohen, L., 2007. Cultural recycling of cortical maps. Neuron, 56(2), pp.384–398.

19. Dehaene, S., Cohen, L., Morais, J. and Kolinsky, R., 2015. Illiterate to literate: behavioural and cerebral changes induced by reading acquisition. Nature Reviews Neuroscience, 16(4), pp.234–244.

20. DiCarlo, J.J. and Cox, D.D., 2007. Untangling invariant object recognition. Trends in cognitive sciences, 11(8), pp.333–341.

21. Dobs, K., Martinez, J., Kell, A.J. and Kanwisher, N., 2022. Brain-like functional specialization emerges spontaneously in deep neural networks. Science advances, 8(11), p.eabl8913.

22. Downing, P.E., Chan, A.Y., Peelen, M.V., Dodds, C.M. and Kanwisher, N., 2006. Domain specificity in visual cortex. Cerebral cortex, 16(10), pp.1453–1461.

23. Downing, P.E., Jiang, Y., Shuman, M. and Kanwisher, N., 2001. A cortical area selective for visual processing of the human body. Science, 293(5539), pp.2470–2473.

24. Durbin, R. and Mitchison, G., 1990. A dimension reduction framework for understanding cortical maps. Nature, 343(6259), pp.644–647.

25. Epstein, R. and Kanwisher, N., 1998. A cortical representation of the local visual environment. Nature, 392(6676), pp.598–601.

26. Freeman, J. and Simoncelli, E.P., 2011. Metamers of the ventral stream. Nature neuroscience, 14(9), pp.1195–1201.

27. Freiwald, W.A. and Tsao, D.Y., 2010. Functional compartmentalization and viewpoint generalization within the macaque face-processing system. Science, 330(6005), pp.845–851.

28. Geirhos, R., Rubisch, P., Michaelis, C., Bethge, M., Wichmann, F.A. and Brendel, W., 2018. ImageNet-trained CNNs are biased towards texture; increasing shape bias improves accuracy and robustness. arXiv preprint arXiv:1811.12231.

29. Graziano, M.S. and Aflalo, T.N., 2007. Mapping behavioral repertoire onto the cortex. Neuron, 56(2), pp.239–251.

30. Grill-Spector, K. and Weiner, K.S., 2014. The functional architecture of the ventral temporal cortex and its role in categorization. Nature Reviews Neuroscience, 15(8), pp.536–548.

31. Hasson, U., Harel, M., Levy, I. and Malach, R., 2003. Large-scale mirror-symmetry organization of human occipito-temporal object areas. Neuron, 37(6), pp.1027–1041.

32. Haxby, J.V., Gobbini, M.I., Furey, M.L., Ishai, A., Schouten, J.L. and Pietrini, P., 2001. Distributed and overlapping representations of faces and objects in ventral temporal cortex. Science, 293(5539), pp.2425–2430.

33. Haxby, J.V., Guntupalli, J.S., Connolly, A.C., Halchenko, Y.O., Conroy, B.R., Gobbini, M.I., Hanke, M. and Ramadge, P.J., 2011. A common, high-dimensional model of the representational space in human ventral temporal cortex. Neuron, 72(2), pp.404–416.

34. Hebart, M.N., Contier, O., Teichmann, L., Rockter, A., Zheng, C.Y., Kidder, A., Corriveau, A., Vaziri-Pashkam, M. and Baker, C.I., 2022. THINGS-data: A multimodal collection of large-scale datasets for investigating object representations in brain and behavior. bioRxiv.

35. Hebart, M.N., Dickter, A.H., Kidder, A., Kwok, W.Y., Corriveau, A., Van Wicklin, C. and Baker, C.I., 2019. THINGS: A database of 1,854 object concepts and more than 26,000 naturalistic object images. PloS one, 14(10), p.e0223792.

36. Hermann, K., Chen, T. and Kornblith, S., 2020. The origins and prevalence of texture bias in convolutional neural networks. Advances in Neural Information Processing Systems, 33, pp.19000–19015.

37. Huang, T., Song, Y. and Liu, J., 2022. Real-world size of objects serves as an axis of object space. Communications biology, 5(1), pp.1–12.

38. Ishai, A., Ungerleider, L.G., Martin, A., Schouten, J.L. and Haxby, J.V., 1999. Distributed representation of objects in the human ventral visual pathway. Proceedings of the National Academy of Sciences, 96(16), pp.9379–9384.

39. Jagadeesh, A.V. and Gardner, J.L., 2022. Texture-like representation of objects in human visual cortex. Proceedings of the National Academy of Sciences, 119(17), p.e2115302119.

40. Julian, J.B., Ryan, J. and Epstein, R.A., 2017. Coding of object size and object category in human visual cortex. Cerebral cortex, 27(6), pp.3095–3109.

41. Kanwisher, N., 2010. Functional specificity in the human brain: a window into the functional architecture of the mind. Proceedings of the National Academy of Sciences, 107(25), pp.11163–11170.

42. Kanwisher, N., McDermott, J. and Chun, M.M., 1997. The fusiform face area: a module in human extrastriate cortex specialized for face perception. Journal of neuroscience, 17(11), pp.4302–4311.

43. Keller, T.A., Gao, Q. and Welling, M., 2021. Modeling category-selective cortical regions with topographic variational autoencoders. arXiv preprint arXiv:2110.13911.

44. Khosla, M. and Wehbe, L., 2022. High-level visual areas act like domain-general filters with strong selectivity and functional specialization. bioRxiv.

45. Kohonen, T., 1990. The self-organizing map. Proceedings of the IEEE, 78(9), pp.1464–1480.

46. Konkle, T. and Alvarez, G.A., 2022. A self-supervised domain-general learning framework for human ventral stream representation. Nature communications, 13(1), pp.1–12.

47. Konkle, T. and Caramazza, A., 2013. Tripartite organization of the ventral stream by animacy and object size. Journal of Neuroscience, 33(25), pp.10235–10242.

48. Konkle, T. and Caramazza, A., 2017. The large-scale organization of object-responsive cortex is reflected in resting-state network architecture. Cerebral cortex, 27(10), pp.4933–4945.

49. Konkle, T. and Oliva, A., 2012. A real-world size organization of object responses in occipitotemporal cortex. Neuron, 74(6), pp.1114–1124.

50. Konkle, T., 2021. Emergent organization of multiple visuotopic maps without a feature hierarchy. bioRxiv.

51. Kourtzi, Z. and Connor, C.E., 2011. Neural representations for object perception: structure, category, and adaptive coding. Annual review of neuroscience, 34, pp.45–67.

52. Kravitz, D.J., Saleem, K.S., Baker, C.I., Ungerleider, L.G. and Mishkin, M., 2013. The ventral visual pathway: an expanded neural framework for the processing of object quality. Trends in cognitive sciences, 17(1), pp.26–49.

53. Kriegeskorte, N., Mur, M., Ruff, D.A., Kiani, R., Bodurka, J., Esteky, H., Tanaka, K. and Bandettini, P.A., 2008. Matching categorical object representations in inferior temporal cortex of man and monkey. Neuron, 60(6), pp.1126–1141.

54. Krizhevsky, A.; Sutskever, I. & Hinton, G. E. (2012), ImageNet Classification with Deep Convolutional Neural Networks, in F. Pereira; C. J. C. Burges; L. Bottou & K. Q. Weinberger, ed., ‘Advances in Neural Information Processing Systems 25’, Curran Associates, Inc.,, pp. 1097--1105

55. Lee, H., Margalit, E., Jozwik, K.M., Cohen, M.A., Kanwisher, N., Yamins, D.L. and DiCarlo, J.J., 2020. Topographic deep artificial neural networks reproduce the hallmarks of the primate inferior temporal cortex face processing network. bioRxiv.

56. Li, S.P.D. and Bonner, M., 2020, October. Curvature as an Organizing Principle of Mid-level Visual Representation: A Semantic-preference Mapping Approach. In NeurIPS 2020 Workshop SVRHM.

57. Long, B., Yu, C.P. and Konkle, T., 2018. Mid-level visual features underlie the high-level categorical organization of the ventral stream. Proceedings of the National Academy of Sciences, 115(38), pp.E9015–E9024.

58. Mahon, B.Z. and Caramazza, A., 2011. What drives the organization of object knowledge in the brain?. Trends in cognitive sciences, 15(3), pp.97–103.

59. Malach, R., Levy, I. and Hasson, U., 2002. The topography of high-order human object areas. Trends in cognitive sciences, 6(4), pp.176–184.

60. Martin, A., 2007. The representation of object concepts in the brain. Annual review of psychology, 58, p.25.

61. Martin, A., Wiggs, C.L., Ungerleider, L.G. and Haxby, J.V., 1996. Neural correlates of category-specific knowledge. Nature, 379(6566), pp.649–652.

62. McCandliss, B.D., Cohen, L. and Dehaene, S., 2003. The visual word form area: expertise for reading in the fusiform gyrus. Trends in cognitive sciences, 7(7), pp.293–299.

63. McCarthy, G., Puce, A., Gore, J.C. and Allison, T., 1997. Face-specific processing in the human fusiform gyrus. Journal of cognitive neuroscience, 9(5), pp.605–610.

64. Mehrer, J., Spoerer, C.J., Jones, E.C., Kriegeskorte, N. and Kietzmann, T.C., 2021. An ecologically motivated image dataset for deep learning yields better models of human vision. Proceedings of the National Academy of Sciences, 118(8), p.e2011417118.

65. Naselaris, T., Stansbury, D.E. and Gallant, J.L., 2012. Cortical representation of animate and inanimate objects in complex natural scenes. Journal of Physiology-Paris, 106(5-6), pp.239–249.

66. Nasr, S., Liu, N., Devaney, K.J., Yue, X., Rajimehr, R., Ungerleider, L.G. and Tootell, R.B., 2011. Scene-selective cortical regions in human and nonhuman primates. Journal of Neuroscience, 31(39), pp.13771–13785.

67. Obermayer, K., Ritter, H. and Schulten, K., 1990. A principle for the formation of the spatial structure of cortical feature maps. Proceedings of the National Academy of Sciences, 87(21), pp.8345–8349.

68. Olah, C., Mordvintsev, A. and Schubert, L., 2017. Feature visualization. Distill, 2(11), p.e7.

69. Op de Beeck, H.P., Haushofer, J. and Kanwisher, N.G., 2008. Interpreting fMRI data: maps, modules and dimensions. Nature Reviews Neuroscience, 9(2), pp.123–135.

70. Osher, D.E., Saxe, R.R., Koldewyn, K., Gabrieli, J.D., Kanwisher, N. and Saygin, Z.M., 2016. Structural connectivity fingerprints predict cortical selectivity for multiple visual categories across cortex. Cerebral cortex, 26(4), pp.1668–1683.

71. Paszke, A., Gross, S., Massa, F., Lerer, A., Bradbury, J., Chanan, G., Killeen, T., Lin, Z., Gimelshein, N., Antiga, L. and Desmaison, A., 2019. Pytorch: An imperative style, high-performance deep learning library. Advances in neural information processing systems, 32.

72. Peelen, M.V. and Downing, P.E., 2005. Selectivity for the human body in the fusiform gyrus. Journal of neurophysiology, 93(1), pp.603–608.

73. Peelen, M.V., Bracci, S., Lu, X., He, C., Caramazza, A. and Bi, Y., 2013. Tool selectivity in left occipitotemporal cortex develops without vision. Journal of cognitive neuroscience, 25(8), pp.1225–1234.

74. Peelen, M.V., He, C., Han, Z., Caramazza, A. and Bi, Y., 2014. Nonvisual and visual object shape representations in occipitotemporal cortex: evidence from congenitally blind and sighted adults. Journal of Neuroscience, 34(1), pp.163–170.

75. Plaut, D.C. and Behrmann, M., 2011. Complementary neural representations for faces and words: A computational exploration. Cognitive neuropsychology, 28(3-4), pp.251–275.

76. Polk, T.A., Stallcup, M., Aguirre, G.K., Alsop, D.C., D’esposito, M., Detre, J.A. and Farah, M.J., 2002. Neural specialization for letter recognition. Journal of Cognitive Neuroscience, 14(2), pp.145–159.

77. Prince, J.S. and Konkle, T., 2020. Computational evidence for integrated rather than specialized feature tuning in category-selective regions. Journal of Vision, 20(11), pp.1577–1577.

78. Puce, A., Allison, T., Asgari, M., Gore, J.C. and McCarthy, G., 1996. Differential sensitivity of human visual cortex to faces, letterstrings, and textures: a functional magnetic resonance imaging study. Journal of neuroscience, 16(16), pp.5205–5215.

79. Ratan Murty, N.A., Bashivan, P., Abate, A., DiCarlo, J.J. and Kanwisher, N., 2021. Computational models of category-selective brain regions enable high-throughput tests of selectivity. Nature communications, 12(1), pp.1–14.

80. Russakovsky, O., Deng, J., Su, H., Krause, J., Satheesh, S., Ma, S., Huang, Z., Karpathy, A., Khosla, A., Bernstein, M. and Berg, A.C., 2015. Imagenet large scale visual recognition challenge. International journal of computer vision, 115(3), pp.211–252.

81. Saygin, Z.M., Osher, D.E., Koldewyn, K., Reynolds, G., Gabrieli, J.D. and Saxe, R.R., 2012. Anatomical connectivity patterns predict face selectivity in the fusiform gyrus. Nature neuroscience, 15(2), pp.321–327.

82. Saygin, Z.M., Osher, D.E., Norton, E.S., Youssoufian, D.A., Beach, S.D., Feather, J., Gaab, N., Gabrieli, J.D. and Kanwisher, N., 2016. Connectivity precedes function in the development of the visual word form area. Nature neuroscience, 19(9), pp.1250–1255.

83. Sha, L., Haxby, J.V., Abdi, H., Guntupalli, J.S., Oosterhof, N.N., Halchenko, Y.O. and Connolly, A.C., 2015. The animacy continuum in the human ventral vision pathway. Journal of cognitive neuroscience, 27(4), pp.665–678.

84. Striem-Amit, E. and Amedi, A., 2014. Visual cortex extrastriate body-selective area activation in congenitally blind people “seeing” by using sounds. Current Biology, 24(6), pp.687–692.

85. Striem-Amit, E., Cohen, L., Dehaene, S. and Amedi, A., 2012. Reading with sounds: sensory substitution selectively activates the visual word form area in the blind. Neuron, 76(3), pp.640–652.

86. Taylor, J.C. and Downing, P.E., 2011. Division of labor between lateral and ventral extrastriate representations of faces, bodies, and objects. Journal of Cognitive Neuroscience, 23(12), pp.4122–4137.

87. Tsao, D.Y., Freiwald, W.A., Tootell, R.B. and Livingstone, M.S., 2006. A cortical region consisting entirely of face-selective cells. Science, 311(5761), pp.670–674.

88. Ungerleider, L.G. and Bell, A.H., 2011. Uncovering the visual “alphabet”: advances in our understanding of object perception. Vision research, 51(7), pp.782–799.

89. Ungerleider, L.G., 1982. Two cortical visual systems. Analysis of visual behavior, pp.549–586.

90. Vinken, K., Konkle, T. and Livingstone, M., 2022. The neural code for’face cells’ is not face specific. bioRxiv.

91. Weiner, K.S. and Grill-Spector, K., 2010. Sparsely-distributed organization of face and limb activations in human ventral temporal cortex. Neuroimage, 52(4), pp.1559–1573.

92. Weiner, K.S. and Grill-Spector, K., 2013. Neural representations of faces and limbs neighbor in human high-level visual cortex: evidence for a new organization principle. Psychological research, 77(1), pp.74–97.

93. Zeki, S.M., 1978. Functional specialisation in the visual cortex of the rhesus monkey. Nature, 274(5670), pp.423–428

94. Zhang, Y., Zhou, K., Bao, P. and Liu, J., 2021. Principles governing the topological organization of object selectivities in ventral temporal cortex. bioRxiv.

