## Supplementary File for "Visual object topographic motifs emerge from self-organization of a unified representational space"

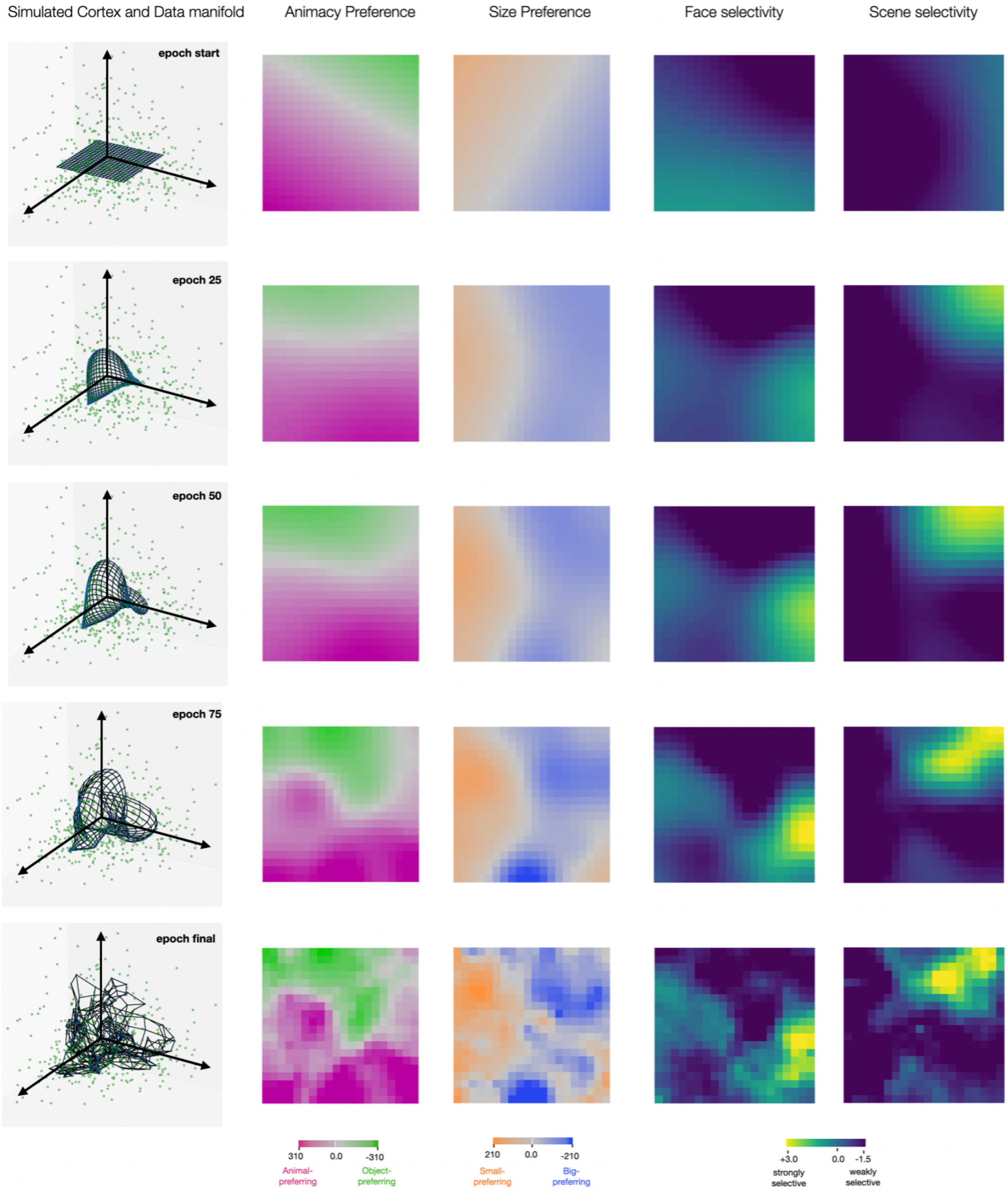


**Supplementary Figure 1.** *Training (Initialization and Fine-tuning) stages of the SOM. In each row we visualize the simulated cortex in context of the input data's PC-space – the green points depict the location of images in the input feature space (dnn features) and the black connected points depict the tuning of SOM map units in this PC-space. We also visualize the animacy and size preference, and face- and scene-selectivity on this simulated cortex for every stage in the training process.*

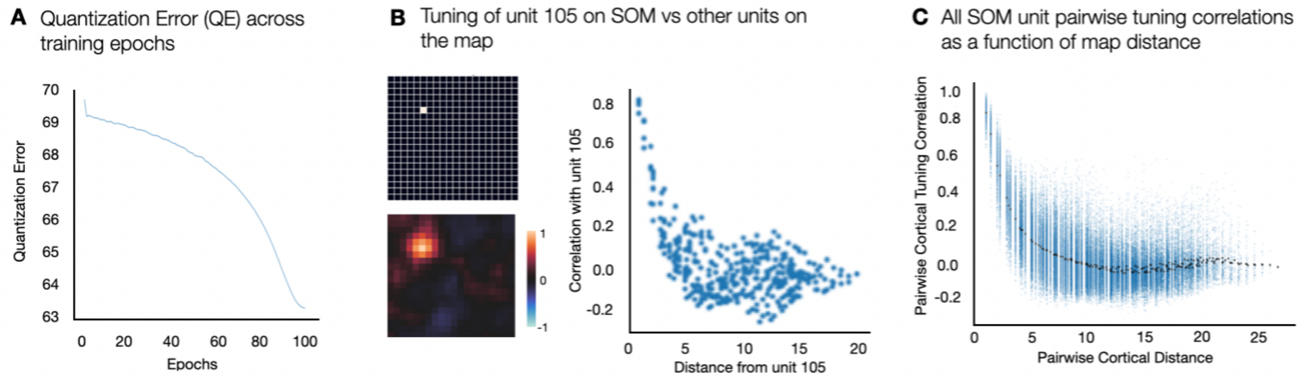


**Supplementary Figure 2.** *(A) Quantization error of the SOM as a function of training epochs (B) Pairwise tuning similarity between one example SOM unit with all other units on the SOM, plotted as a heat map (top) and a scatter plot (below), with tuning similarity (correlation) along the y-axis, plotted as a function of map distance between units (euclidean) on the x-axis (C) Scatter plot with tuning similarity between every pair of units on the SOM on the y-axis and the map distance (euclidean) between pair of units on the x-axis.*

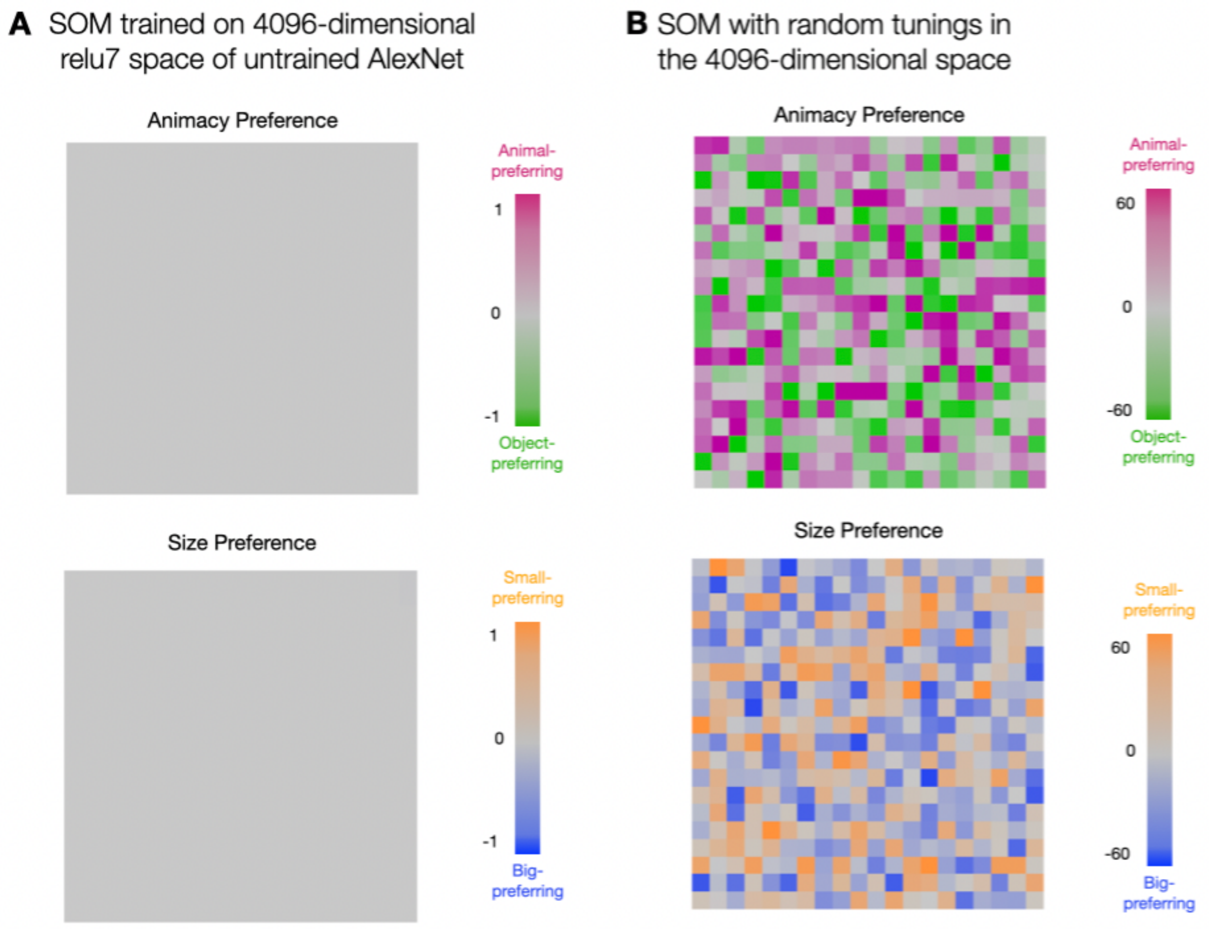


**Supplementary Figure 3.** (A) *Animacy (animals vs objects) and Size (big vs small entities) preference on the simulated cortex over the relu7 ($\mathbb{R}^{4096}$) feature space of an untrained Alexnet. (B) Animacy and size preference on the simulated cortex when the SOM is randomly tuned in a 4096-dimensional space.*

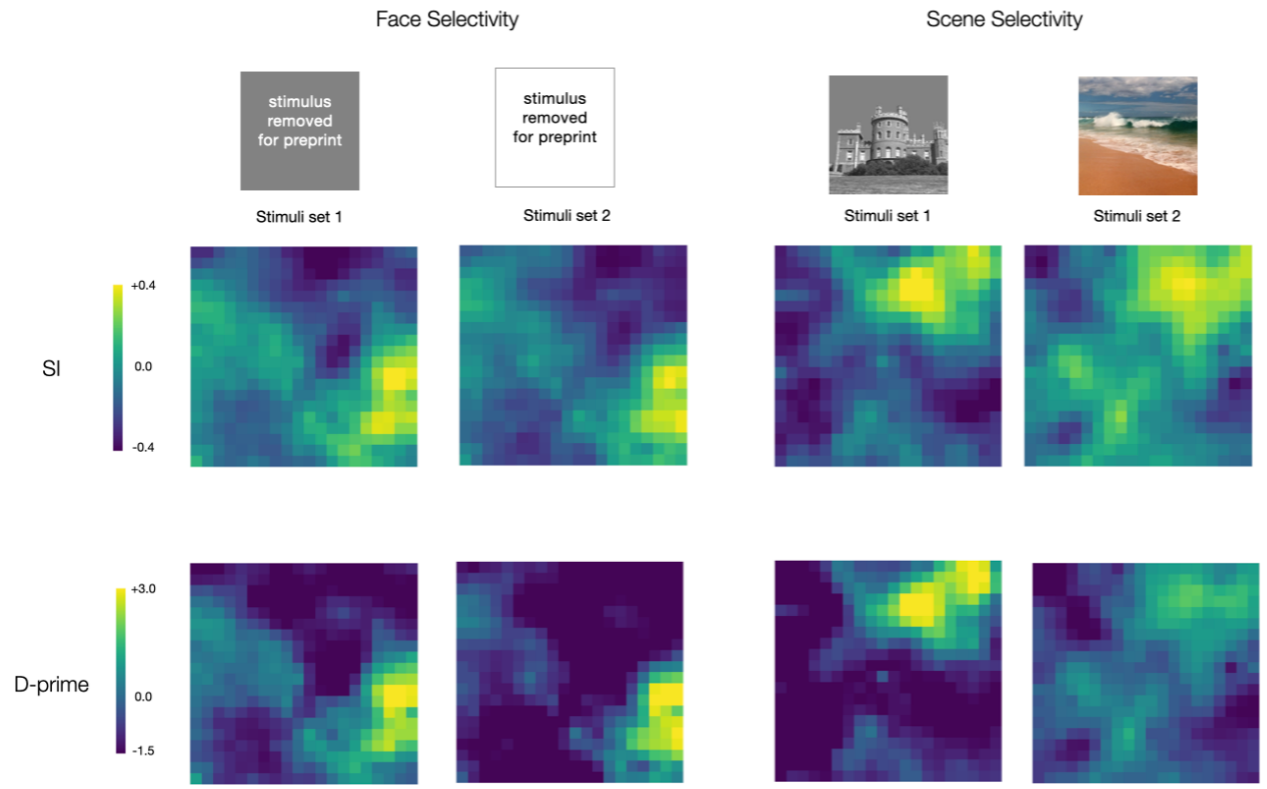


**Supplementary Figure 4.** *Category selectivity for faces and scenes within the two stimuli sets, measured using Selectivity Index (SI) and D-prime measure.*

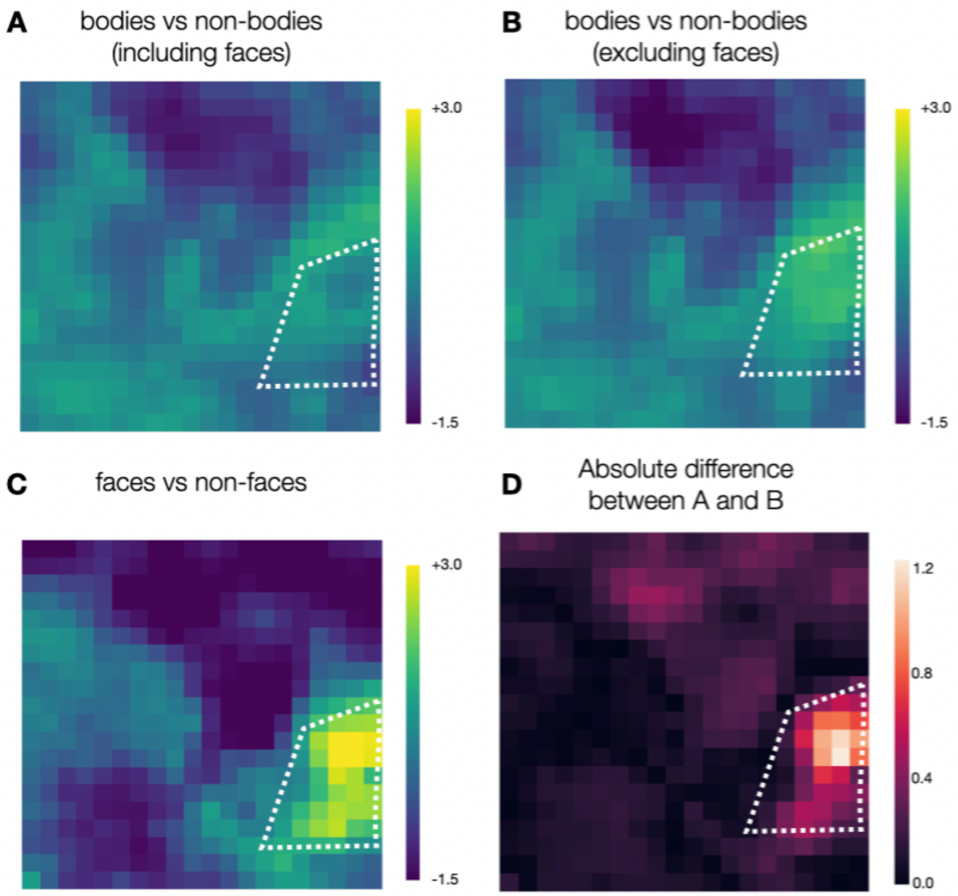


**Supplementary Figure 5.** *(A and B) Body-selectivity on the simulated cortex with faces included and excluded while computing the d-prime selectivity map. (C) Face-selectivity on the simulated cortex (D) Mean Absolute difference between (A) and (B). The white lines demarcate the most face-selective zone.*

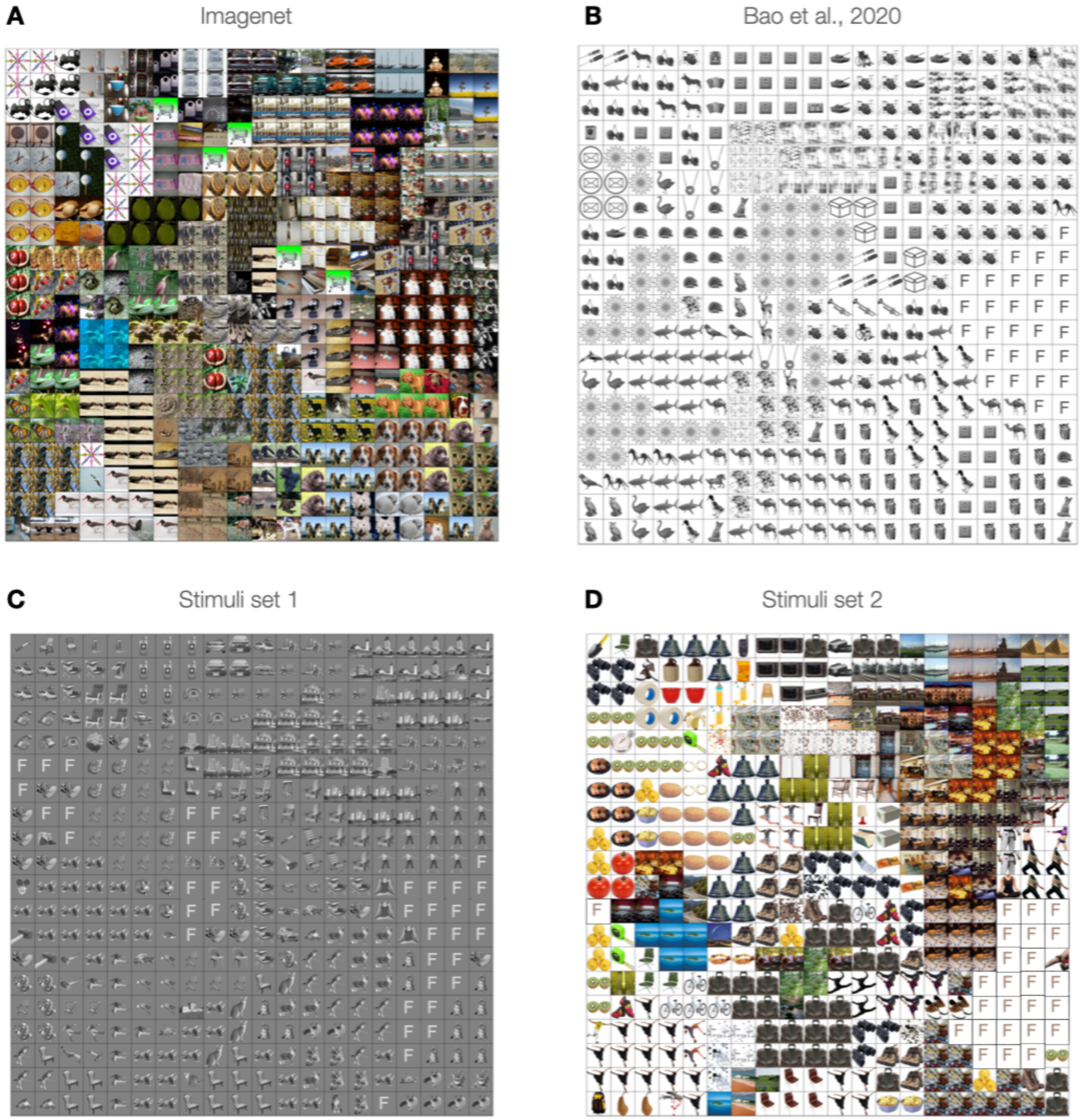


**Supplementary Figure 6.** *Map of images that maximally drive map units on the 20*20 grid computed over different stimulus sets: (A) ImageNet validation set (B) stimuli used in Bao et al., 2020 (C,D) two localizer sets used in this study. Face images replaced with letter F.*

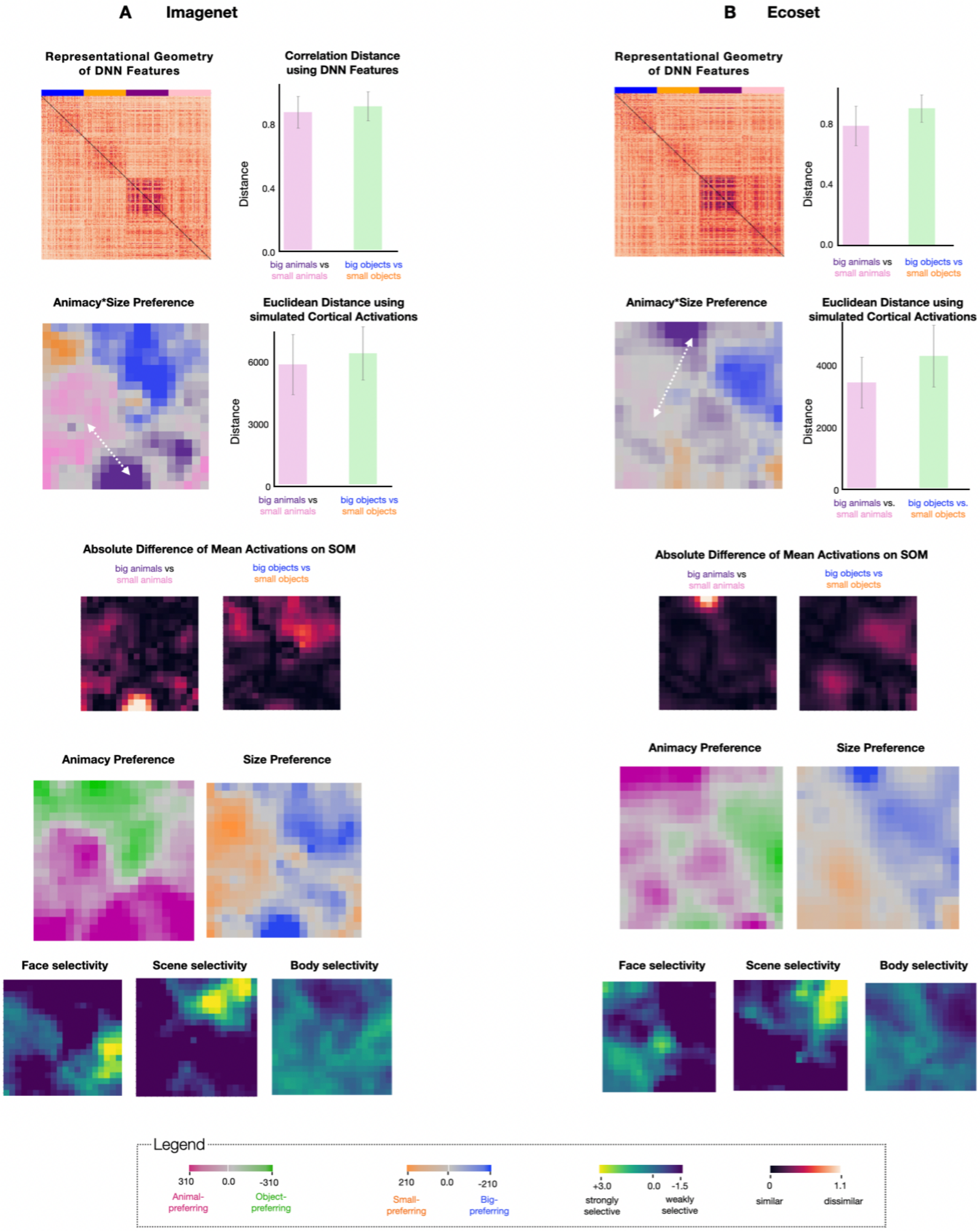


**Supplementary Figure 7.** *Comparing the large-scale organization between models (deepnet and SOM) trained on different visual diets – imagenet vs ecoset. For each visual experience, we visualize: (i) Representational Geometry of images (stimuli from Konkle & Caramazza, 2013) in the deep net feature space. (Left) RDMs based on correlational distance. (Right) Bar plots showing the*

*image-level pairwise correlational distance between dnn features of  big and small animals, and dnn features of big and small objects. (ii) (Left) 4-way preference map on the simulated cortex among big objects, small objects, big animals, and small animals and (Right) ) Bar plots showing the image-level pairwise euclidean distance between simulated cortical activations of big and small animals, and simulated cortical activations of big and small objects*
*using the same stimuli as in (i). (iii) Heatmaps showing the absolute difference of mean simulated cortical activations based on size for animals (i.e. big animals vs. small animals) and objects (i.e. big objects vs. small objects) (iii) Animacy (animals vs. objects) and Size (small vs. big) preferences on the simulated cortex. (iv) Face-, Scene-, and Body-selectivity on the simulated cortex measured using the d-prime measure. Stimulus from Cohen et al., 2017 were used to compute the selectivity maps.*

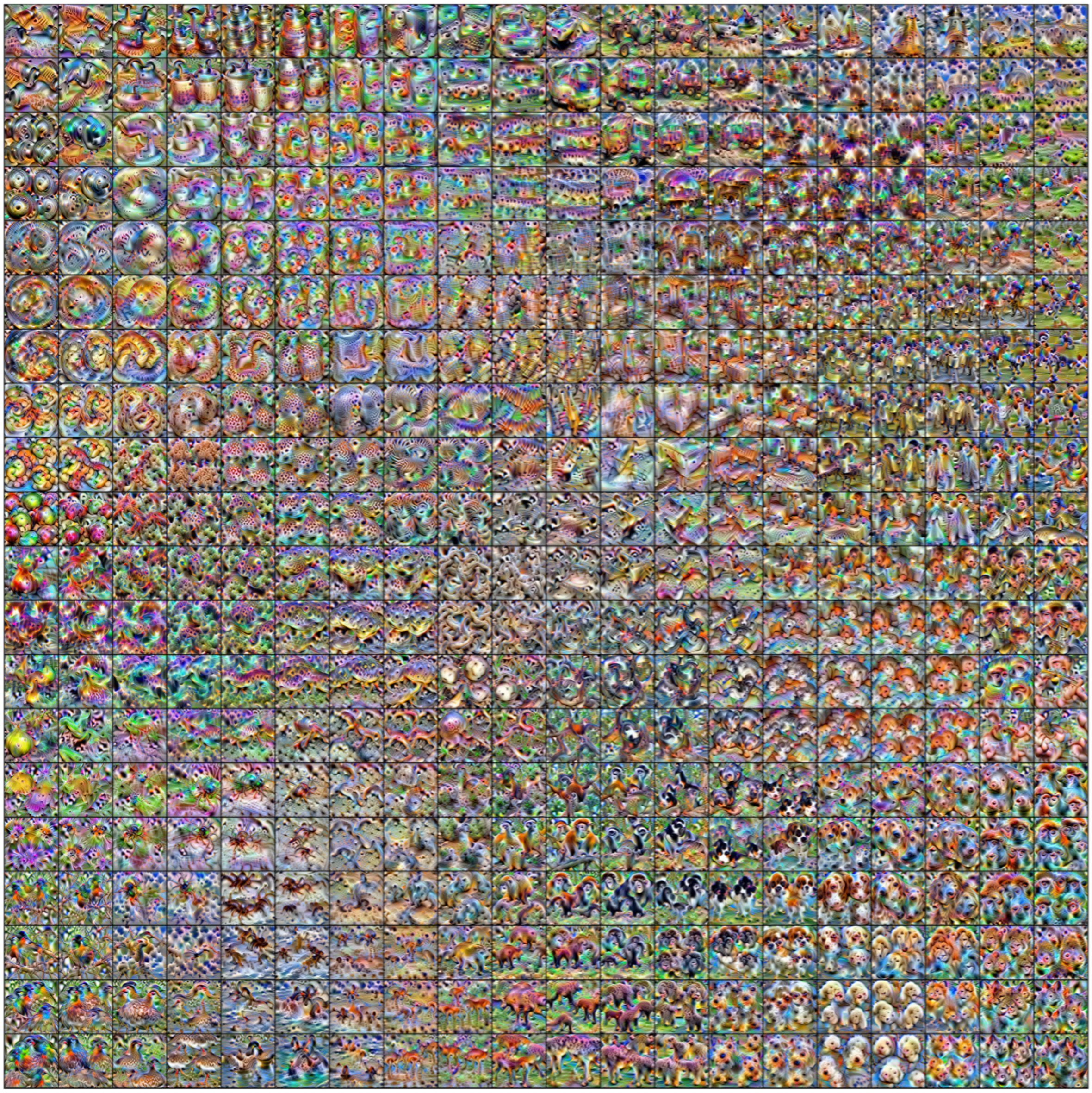

**Supplementary Figure 8.** *Synthesized maximally activating images, generated using gradient ascent, for all map units on the 20*20 grid*

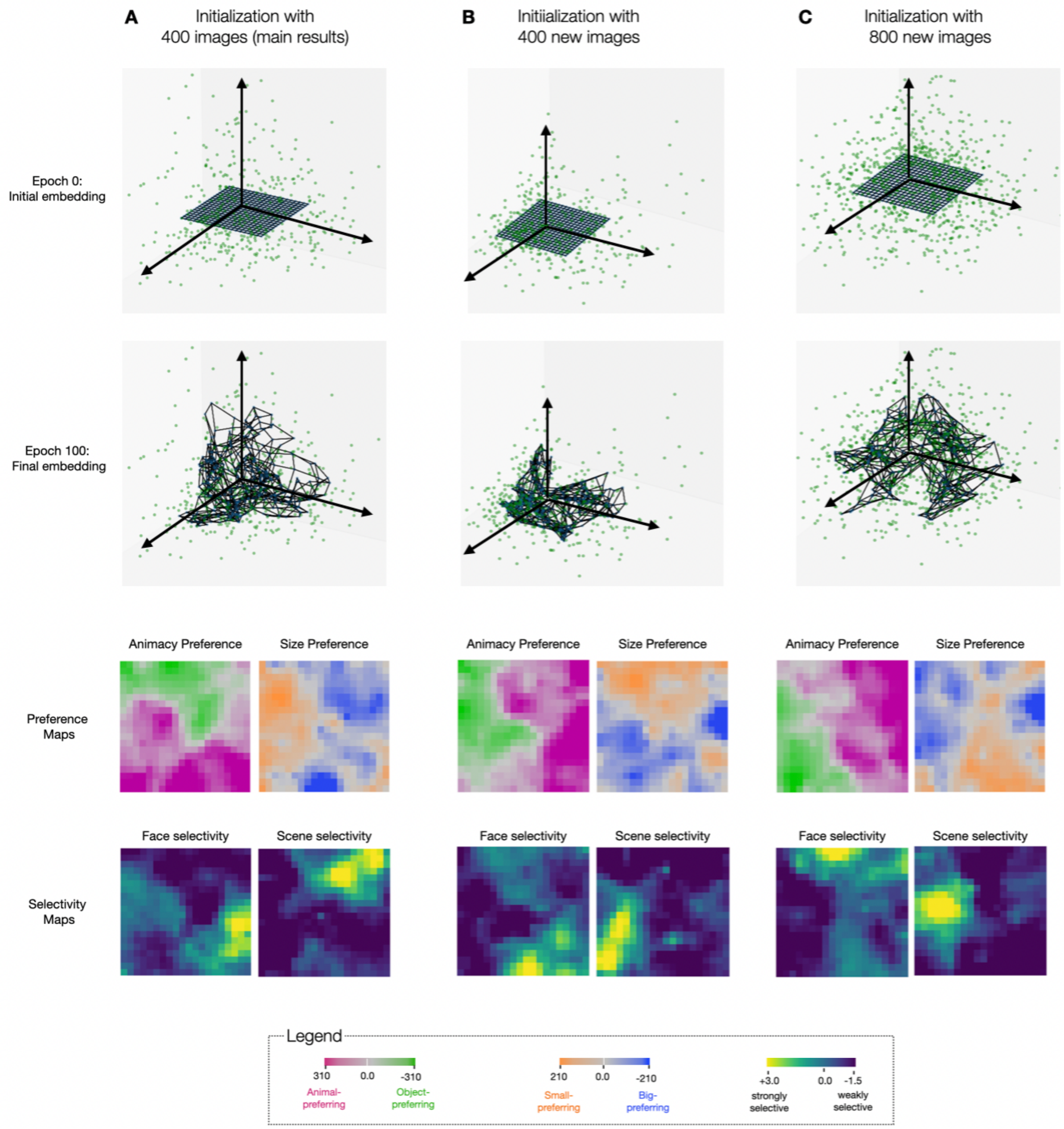


**Supplementary Figure 9.** *Each column is a trained map initialized with different images from the SRS/Imagenet validation set. First column uses 400 images (same as the ones reported in the main figures), the second column uses 400 different images, and the third column uses 800 images. The training images remain the same for all 3 variations. Top and Second row: Visualization of the map units projected into the first three principal components of the input space before (epoch 0) and after training (epoch 100). Third row: Large-scale organization of animacy and size after training. Bottom row: Category selectivity for faces and scenes after training*

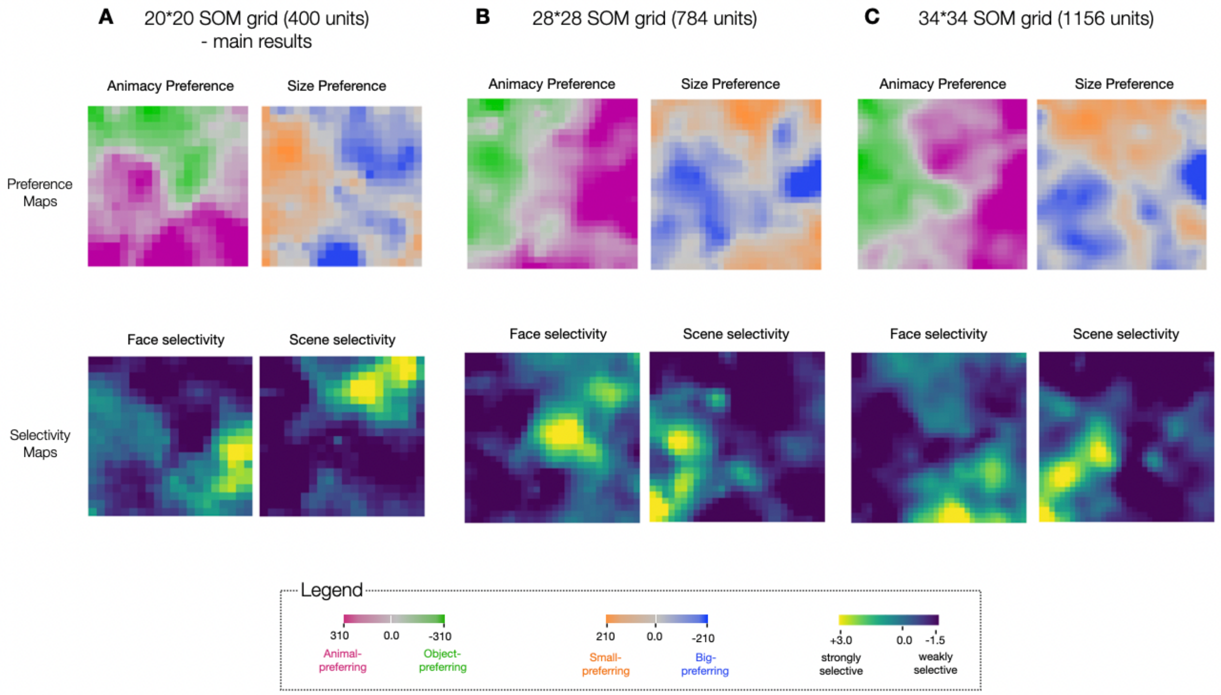


**Supplementary Figure 10.** *Each column is a trained map. The size of the map varies from small to large, from left to right. Top row: Large-scale organization of animacy and size. Bottom row: Category selectivity for faces and scenes*
